# Non-invasive forecasting of skin cancer evolution through longitudinal hair sampling

**DOI:** 10.64898/2026.08.31.748188

**Authors:** Stefania Del Prete, Yoav Avi-Guy, Fu Xu, Sabrina Weser, Milica Bekavac, Marie-Luise Koch, Francesca Coraggio, Raphael T. F. Coimbra, Martyna C. Popis, Anke Heit-Mondrzyk, Angela T. Goncalves, Mikaela Behm, Duncan T. Odom, Michaela Frye

**Author notes:** Co-corresponding: M.F.; D.T.O.

## Abstract

The ability to longitudinally track clonal evolution non-invasively would transform cancer interception strategies, long before late-stage disease when most cancer genomes are analysed. Here, we demonstrate that repeated hair sampling from the same individual followed by exome sequencing enables tracking of somatic evolution *in vivo* over several months after chemically induced skin carcinogenesis. We found that hair follicles accumulate a higher mutation burden than spatially-matched skin and harbour mutations that spread into surrounding epidermis and persist throughout tumour progression. DNA-damaged follicles enter sustained quiescence that delays replication and repair, creating a reservoir for long-lived mutations. During premalignant progression, carcinogen-associated mutations become enriched as follicular clones expand into adjacent skin. Mutation tracking identified genes that may govern tumour predisposition and initiation, many of which are mutated at high incidence in human cutaneous squamous cell carcinoma cohorts. Hair follicles therefore provide a non-invasive readout to forecast the early development of skin cancer, enabling patient risk stratification.

## Introduction

Preventing skin cancer progression requires defining the earliest molecular and clonal events that precede malignant transformation, after which preventive intervention is less likely to be effective. The premalignant stages of carcinogenesis remain difficult to interrogate because they evolve over extended periods of time and cannot readily be sampled longitudinally. Consequently, our understanding of early tumour evolution has largely relied on endpoint analyses and retrospective reconstruction of clonal history.

The two-stage chemical carcinogenesis model recapitulates stepwise progression from mutagenesis through premalignant papillomas to invasive cutaneous squamous cell carcinoma (SCC), the second most common skin cancer ^1^, providing a unique framework to study early carcinogenesis ^2^. In this model, tumour initiation is achieved by introduction of DNA mutations using a single dose of 7,12-dimethylbenz[a]-anthracene (DMBA), followed by stimulated epidermal proliferation via repeated treatment with tetradecanoyl-phorbol acetate (TPA) ^2^. The protocol induces the same spectrum of genomic alterations found in humans ^3^, including activating mutations in *Hras* ^4–6^, a driver frequently found in human cutaneous SCC ^7^.

The prolonged premalignant phase of the DMBA/TPA model enables prospective tracking of somatic mutations, providing direct insight into clonal selection and evolution from tumour initiation through malignant transformation. Hair follicles are uniquely suited for longitudinal studies of skin carcinogenesis because they contain long-lived epithelial stem cells that remain anatomically stable and are readily accessible for repeated sampling throughout life ^8,9^. Recent lineage-tracing studies have identified hair follicle stem cells as an important cellular origin of chemically induced skin tumours and demonstrated the long-term persistence of their progeny during carcinogenesis ^10,11^. These properties suggested to us that hair follicles could serve as a reservoir of persistent mutations while retaining a genomic footprint of neighbouring epidermal evolution, providing an opportunity to monitor the earliest stages of carcinogenesis before overt tumour formation.

Here, we developed a non-invasive longitudinal strategy that enables repeated, spatially matched sampling of hair follicles from individual mice throughout multistage skin carcinogenesis. We reasoned that hair follicles may serve as genomic sentinels of neighbouring epidermal evolution, allowing prospective tracking of clonal trajectories from tumour initiation to malignant transformation. This approach provides a foundation for identifying the earliest genomic determinants of tumour initiation and regional tumour predisposition, unlocking the ability to intervene well before malignant transformation.

## Results

### Spatiotemporal analysis of somatic mutations in hair follicles during skin carcinogenesis

Although hair follicle stem cells can give rise to cutaneous squamous cell carcinoma (SCC) ^11–14^, it has not previously been possible to track genomic mutations in these cells longitudinally during multistage skin carcinogenesis. Here, we developed a non-invasive tape-stripping method for longitudinal *in vivo* sampling of hair follicles during skin carcinogenesis. Hair follicles were repeatedly sampled from the same mouse undergoing two-stage chemical carcinogenesis (Figure 1A), in which tumour initiation was induced by a single DMBA application followed by repeated TPA-mediated tumour promotion ^2^. All mice developed papillomas within ∼10 weeks of treatment, as expected; in only two animals these progressed to an SCC within the 20-week treatment time window (Figure 1B, C; Supplementary Figure 1A, B).

**Figure 1.**
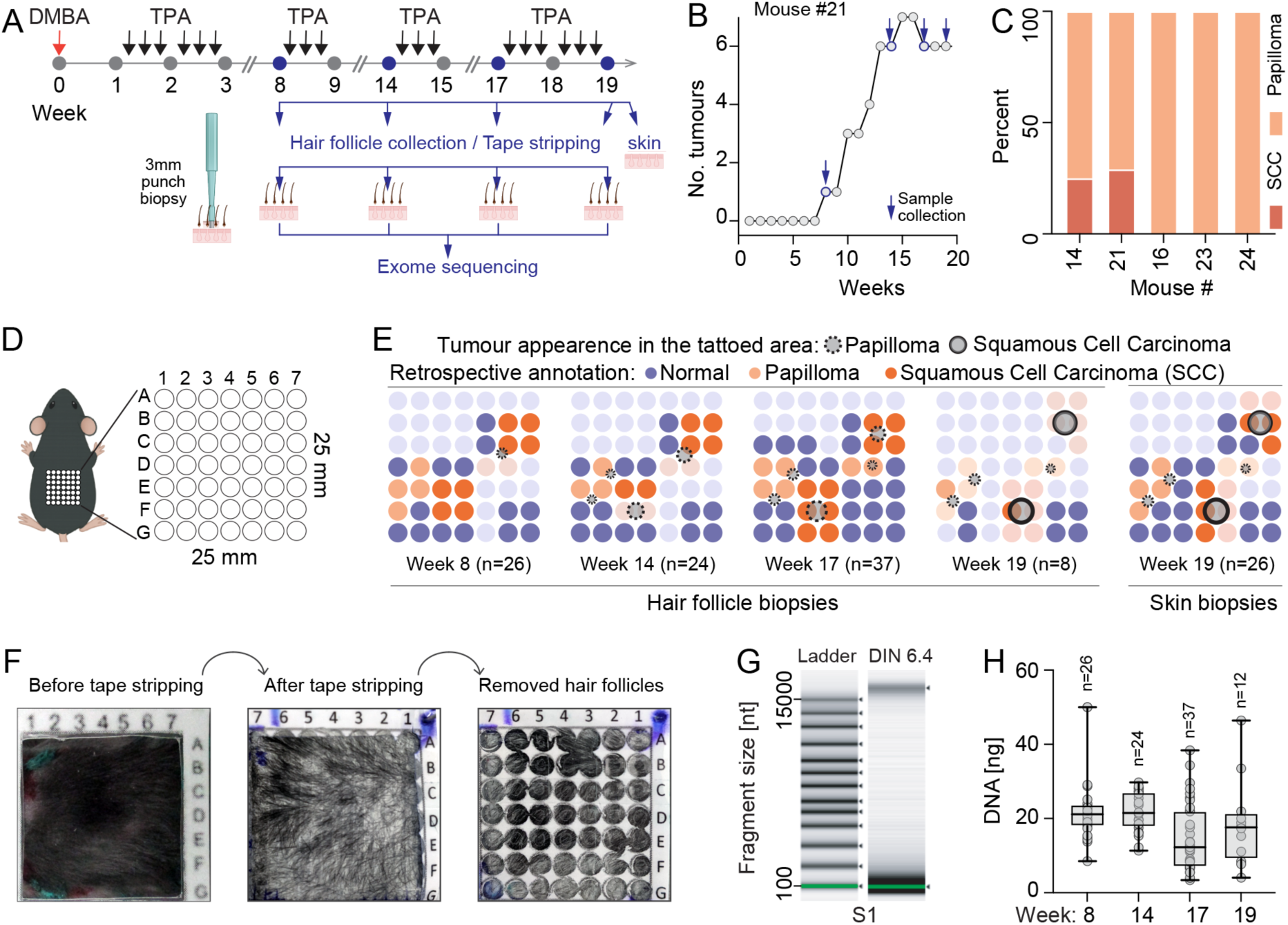
Non-invasive longitudinal sampling for spatially resolved mutational profiling. (**A**) Experimental design of DMBA/TPA-induced skin carcinogenesis and longitudinal sampling strategy. Hair follicles were collected at weeks 8, 14, 17, and 19. Matched skin samples were collected at week 19. (**B, C**) Number of tumours (B) and tumour types (C) in mice used for the chemical carcinogenesis experiment. SCC: Squamous cell carcinoma. (**D, E**) Illustration of the spatially resolved grid used for collection of hair follicles and skin samples (D). Retrospective annotation of areas (E) where SCCs (grey circles with solid line) or papillomas (grey circles with dotted line) occurred over the course of the experiment using as reference the 19-week endpoint (skin biopsies). Dark orange grid dots: punch biopsies around SCCs. Light orange grid dots: punch biopsies around papillomas. Blue grid dots: punch biopsies around normal skin. Pale blue and orange circles: not analysed. (**F-H**) Representative examples of workflow (F), DNA quality (G) and DNA yield of isolated hair follicles at weeks 8, 14, 17, and 19 (H). n: Number of biopsies (E, H). Box plots show minimum, first quartile, median, third quartile, and maximum (H). Icon of punch biopsy blade in the figure panel A was created with BioRender.com.

A standardised tattooed biopsy grid enabled repeated minimally invasive collection of hair follicles in live animals from spatially matched skin regions (Figure 1D, E). Hair follicle biopsy grid positions were retrospectively classified according to terminal skin lesion morphology as: macroscopically normal, papillomas (regressed tumour sites) or SCCs (Figure 1E). Early time point grid positions that ultimately developed papillomas or SCCs are hereafter referred to as tumour-destined regions. Full-thickness skin biopsies collected at the study endpoint served as a tissue reference for comparison with longitudinal hair follicle biopsies (Figure 1A, E). DNA was extracted from hair follicles captured on tape strips and isolated using individual 3-mm punch biopsies (Figure 1F). Hair follicle samples consistently yielded 3 to 50 ng of high quality DNA (Figure 1G, H). DNA extracted from hair follicle and skin biopsies was used in targeted exome sequencing reactions covering 225 SCC-related genes (Supplementary Table1) ^3^ or in whole-exome sequencing reactions to measure high-resolution mutational changes over time (Supplementary Figure 1C-H). Thus, we established a robust, non-invasive methodology that allows longitudinal tracking of somatic mutations *in vivo* during skin carcinogenesis.

### Hair follicles exhibit distinct responses to mutagenic injury

To quantify somatic mutations in skin and hair follicle biopsies, we used matched liver tissue from the same animals as a germline reference to exclude germline variants. Overall, hair follicles harboured a significantly higher mutation burden than matched full-thickness skin biopsies (Figure 2A). This finding raised the surprising possibility that hair follicle cells possess distinct intrinsic properties that allowed the persistence and accumulation of mutations. To determine whether hair follicle cells acutely respond to a chemical mutagen differently than neighbouring interfollicular epidermis (IFE), we investigated the compartment-specific impact at 48 hours after a single topical dose of DMBA. We first asked whether CD34+ hair follicle stem cells located in the bulge ^15^ sustain damage at all, given their quiescent state and deeper position within the skin. Immunofluorescence staining for γH2AX, which marks DNA damage induced by DMBA, revealed a significant increase of nuclear foci in the quiescent bulge, the upper hair follicle region and the IFE (Figure 2B, D). In contrast, the proliferation marker Ki67 was significantly reduced in the upper hair follicle and IFE compartments (Figure 2C, E), indicating that DNA damage was not accompanied by proliferation at this time point.

**Figure 2.**
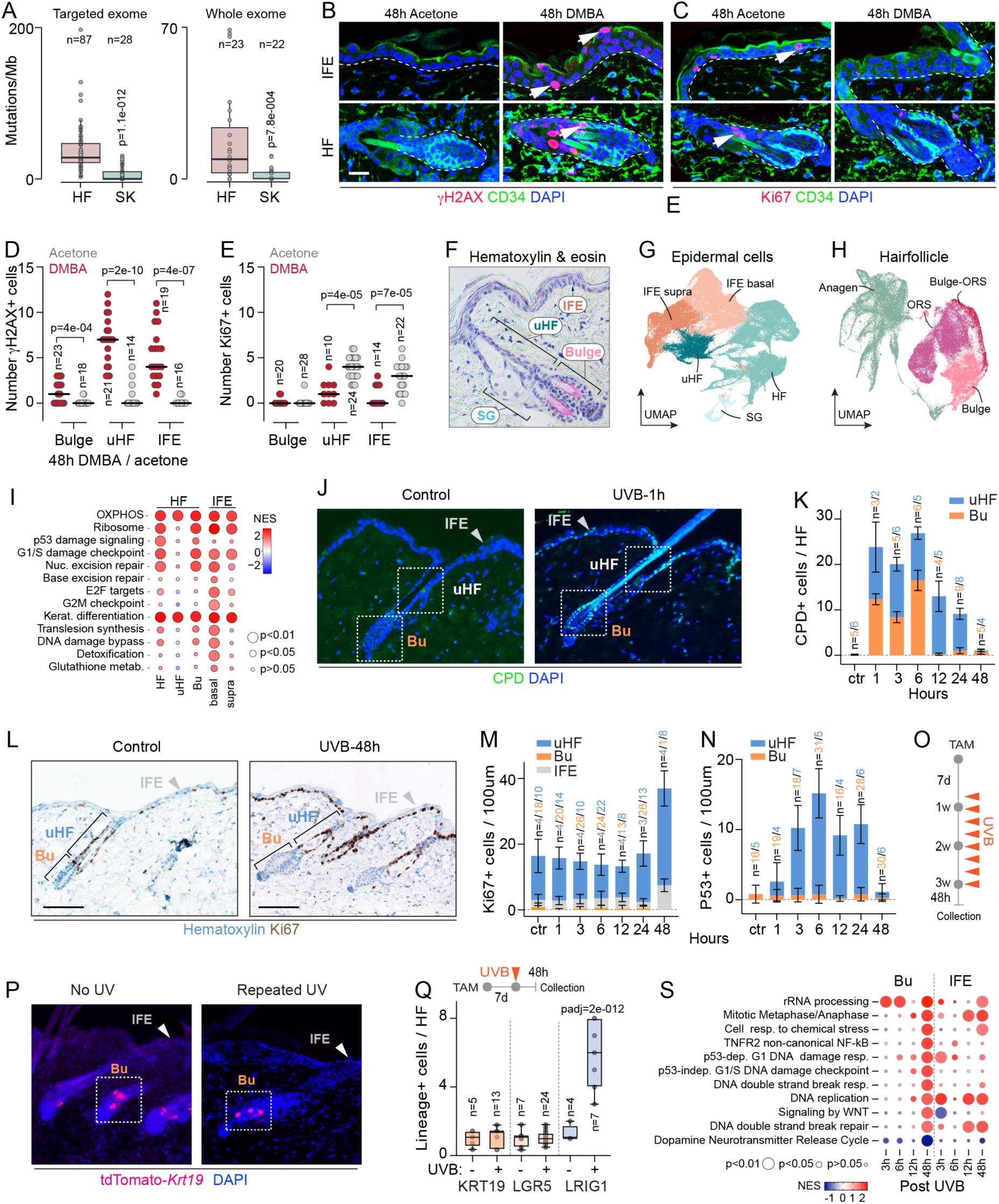
Skin compartment-specific responses to mutagenic stress. (**A**) Somatic mutation burden including nucleotide variants and indels per megabase (Mb) measured by targeted exome (left) and whole exome sequencing (right) in hair follicles (HF) and skin (SK) biopsies after DMBA/TPA treatment. n: Number of biopsies. (**B-E**) Immunofluorescence labelling (B, C) and quantification (D, E) of γH2AX-(B, D) and Ki67-(C, E) positive nuclei in the interfollicular epidermis (IFE; counted as region spanning between two HFs), the upper hair follicle (uHF), and CD34+ bulge cells 48 hours (h) after a single dose of DMBA or acetone used as a control. (**F**) Illustration showing the different epidermal compartments. SG: Sebaceous gland. (**G, H**) Uniform Manifold Approximation and Projection (UMAP) of all epidermal (G) and hair follicle (H) cells. IFE is separated into basal (undifferentiated) (IFE basal) and suprabasal (IFE supra) epidermal cells (G). Hair follicle cells are separated into bulge and outer root sheath (ORS) cells. (**I**) Gene set enrichment analysis of transcriptional changes of interfollicular and follicular sub-compartments. Shown are the top pathways significantly changing and shared between cell types. Colour code represents normalised enrichment scores (NES), size represents adjusted P-values. (**J, K**) Immunofluorescence labelling (J) and quantification (K) of CPD-positive in the uHF and bulge after a single dose of UVB irradiation at the indicated time points. (**L-N**) Immunohistochemistry staining (L) and quantification (M, N) of Ki67-(L, M) and P53-(N) positive nuclei in the IFE and uHF in control or UVB-treated skin. n: Labelled-positive cells per hair follicle (K) or per 100 μm (M, N) obtained from three different mice. (**O, P**) Experimental design of repeated UVB-treatment regimen (O) and detection of tdTomato in untreated (No UV) and UVB-treated (repeated UV) back skin in genetically labelled *Krt19* cell populations. Arrowheads: tdTomato-positive cells. Dotted square: Bulge. (**Q**) Experimental design (top) and quantification of tdTomato-positive cells per hair follicle shown as fold-change over untreated control in the indicated lineage-traced population (bottom). (**S**) Gene ontology analysis of transcriptional changes in bulge and IFE cell populations isolated as shown in Supplementary Figure 2L. Shown are the top pathways by adjusted P-values. Colour code represents normalised enrichment scores (NES), size represents adjusted P-values. n = number of compartments analysed across three different mice (D, E, K, M, N, Q). Median (D, E). Mean ± SD (K, M, N,). Box plots show minimum, first quartile, median, third quartile, and maximum (Q). Kruskal-Wallis test (A). Unpaired t test with Welch’s correction (D, E). Šídák’s multiple comparisons test (Q).

To dissect transcriptional differences between follicular and interfollicular compartments, we performed single-cell RNA sequencing on dissociated whole skin, and then resolved interfollicular and follicular sub-compartments using unsupervised clustering (Fig. 2F-H; Supplementary Figure 2A-D). Gene set enrichment analysis revealed compartment-specific transcriptional responses across epidermal regions (Figure 2I; Supplementary Table 2). However, all compartments sensed the damage, engaging the DNA damage checkpoint and nucleotide excision repair, together with coordinated upregulation of oxidative phosphorylation and ribosomal gene expression. This response was not proliferative, consistent with the reduction of Ki67 immunostaining (Figure 2C, E); enrichment of replication-annotated gene sets was instead driven by histone, proteasome and repair genes (Supplementary Table 2). Epidermal keratinocytes therefore respond to DMBA by arresting and differentiating rather than by proliferating, while simultaneously increasing biosynthetic and respiratory capacity. Although damage sensing was shared, its resolution was compartment-specific. Follicular cells were not repair-deficient, but the bulge showed enrichment of DNA damage bypass, most plausibly reflecting error-prone gap-filling synthesis accompanying excision repair (Figure 2I). Together, these data indicate that bulge stem cells survive DMBA-induced damage in a p53-arrested state, while engaging low-fidelity DNA synthesis. Because bulge stem cells are long-lived and clonally persistent, this provides a mechanistic basis for the enrichment of mutations we observe in hair follicles.

We next investigated whether ultraviolet (UVB) irradiation, a major environmental mutagen and etiological driver of human skin carcinogenesis, elicits compartment-specific responses like those induced by DMBA. A single dose of UVB irradiation was sufficient to generate cyclobutane pyrimidine dimers (CPDs), which cause most UV-induced mutations and can readily be detected by immunostaining (Figure 2J) ^16–18^. CPDs were present in cells located in both the bulge and upper hair follicle as early as one hour post UV-irradiation, yet damaged bulge cells were cleared faster than in the upper hair follicle (Figure 2J, K; Supplementary Figure 2E). Despite this detectable DNA damage, bulge cells showed no up-regulation of proliferation or evidence of apoptosis (Figure 2L-N; Supplementary Figure 2F, G). When we tracked genetically labelled single cells located in the hair follicle bulge (*Krt19; Lgr5*) or the junctional zone of the upper hair follicle (*Lrig1; Lgr6*) (Supplementary Figure 2H), we found that repeated exposure to UVB expanded individual junctional zone cells, whereas bulge-derived cells showed no expansion compared with untreated controls (Figure 2O, P; Supplementary Figure 2I, J). Similarly, only upper hair follicle cells clonally expanded after a single dose of UVB (Figure 2Q; Supplementary Figure 2K). Thus, as observed with DMBA-induced DNA damage, UVB irradiation induced compartment-specific responses.

To identify the underlying molecular mechanisms, we isolated CD34+ bulge stem cells and ITGA6+ undifferentiated interfollicular epidermis (which includes the junctional zone) using flow sorting and performed bulk transcriptomic analysis (Supplementary Figure 2L). Our analysis revealed that both bulge and IFE compartments detected and responded to UV, but resolved the resulting damage differently (Figure 2S; Supplementary Table 3). Bulge cells displayed a relatively delayed response, with the strongest pathway enrichment observed at 48 hours after UVB exposure, including p53-dependent and -independent DNA damage checkpoints, DNA double-strand break responses and repair, DNA replication, and cellular stress pathways. In contrast, IFE cells showed a more dynamic response across time, with early activation of DNA replication and p53-associated pathways followed by prominent enrichment of mitotic and DNA repair programs at later time points. These findings suggest that bulge cells mount a delayed but coordinated DNA damage response to UVB, characterized by restrained post-damage proliferation and DNA replication, whereas IFE cells exhibit a more heterogeneous and less temporally organized transcriptional response. This quiescent state and delayed proliferative response may permit the persistence of unrepaired DNA lesions, which can subsequently become fixed as mutations upon cell-cycle re-entry, potentially contributing to the elevated mutational burden observed in hair follicles.

### Mutational signatures in hair follicles dynamically evolve during tumorigenesis

Mutagen exposure alone is often insufficient for tumour formation; a proliferative stimulus is commonly required for cancer development ^2,19^. We reasoned that hair follicles may serve as a reservoir in which quiescence allows mutations to accumulate to high concentrations specifically in the bulge, which could then be clonally expanded by replicative activation.

To test this, we examined how repeated TPA treatment, which drives anagen entry and transient efflux of bulge-derived cells into other epidermal compartments (Figure 3A) ^20–23^, induces mutational changes in hair follicles over time. Consistent with the anticipated loss of quiescence following TPA treatment, the number of Ki67-positive cells in the bulge significantly increased, while the upper hair follicle exhibited the most pronounced proliferative expansion (Figure 3B, D). In contrast, the extent of DNA damage in response to the TPA treatment was comparable across all epidermal compartments examined (Figure 3C, E).

**Figure 3.**
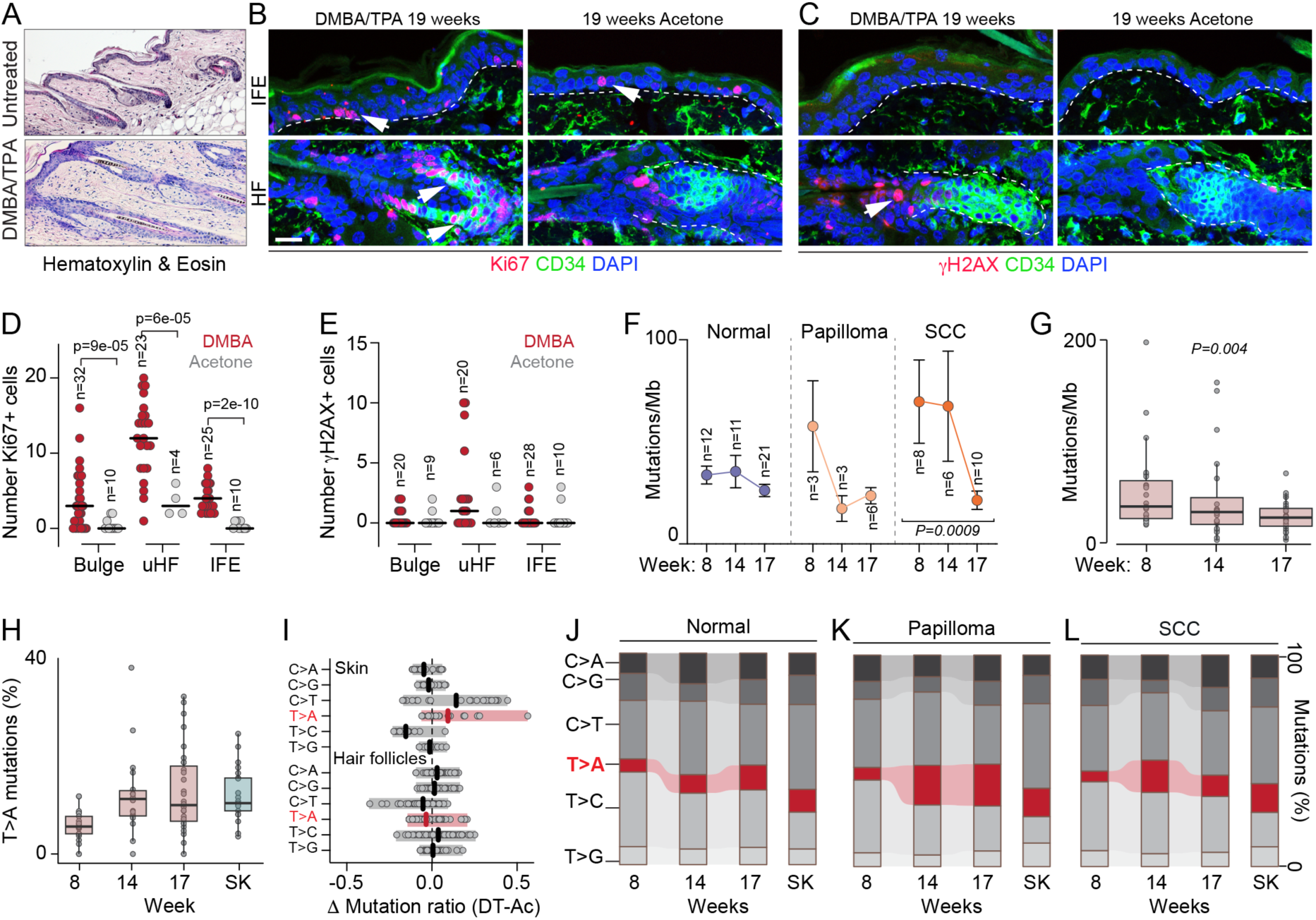
Dynamic changes in mutation burden and signature during tumour development. (**A**) Haematoxylin and Eosin staining of untreated (top) and DMBA/TPA treated (bottom) sections from mouse back skin. (**B-E**) Immunofluorescence labelling (B, C) and quantification (D, E) of Ki67-(B, D) and γH2AX-(C, E) positive nuclei in the interfollicular epidermis (IFE; counted as region spanning between two HFs), the upper hair follicle (uHF), and CD34+ bulge cells after 19 weeks of DMBA/TPA treatment. Acetone treated animals served as controls. n= number of compartments analysed across two DMBA/TPA and one acetone treated mice. (**F, G**) Number of mutations per megabase (Mb) over time after targeted exome sequencing separated by morphological annotation (F) and using all hair follicle biopsies (G). Negative binomial generalised linear models controlling for per-biopsy callable territory were used to test mutation-count differences over time; BH-adjusted P-values are reported. (**H**) Number of T>A DMBA-specific mutations per megabase (Mb) over time in hair follicles and in skin (SK). Points represent individual biopsies; boxplots show the median, interquartile range, and 1.5 IQR whiskers. (**I**) Frequencies of single-nucleotide variants (SNVs) in DMBA/TPA-treated samples following subtraction of SNV contributions from acetone (Ac)-treated samples. (**J-L**) Proportional contribution of SNVs obtained from hair follicles across timepoints (8, 14, 17 weeks) or skin biopsies (19 weeks) in normal (J) or tumour-destined (papilloma, SCC) (K, L) regions. Beta-binomial models adjusting for per-biopsy callable Mb showed a significantly higher T>A proportion in papillomas than in SCC (BH-adjusted P = 0.0046).

As mutant clones expanded, hair follicles in regions from which papillomas or SCCs would later emerge (tumour-destined regions) showed on average the highest mutation burdens by week eight (Figure 3F). Over time the overall mutational burden in the hair follicle decreased (Figure 3G), while the proportion of T>A transversions caused by DMBA exposure increased, and the rate of nonsynonymous mutations remained stable (Figure 3H; Supplementary Figure 3A). This is expected when clonal diversity decreases, due to continuous clonal competition. Overall, within tumour-destined regions, the promotion phase appears to be characterised by clonal evolution associated to both clonal expansion and sweeps (Supplementary Figure 3B).

Across the entire dataset, T>A transversions and C>T transitions were predominant in skin but were not enriched in hair follicles (Figure 3I). Although C>T transitions can arise through several mechanisms, APOBEC cytidine deaminases were transcriptionally repressed in hair follicles when compared to IFE cells, following mutagen exposure (Supplementary Figure 3C), which may contribute to the lack of C>T enrichment in the follicular compartment.

We then determined whether tumour-destined regions showed different longitudinal mutation dynamics than regions that would remain macroscopically normal. C>T transitions were largely unchanged across all conditions, and were generally the most common mutation type (Figure 3J-L). In contrast, T>A transversions increased more rapidly in tumour-destined regions, particularly for papillomas (Figure 3J-L; Supplementary Figure 3D). The greater papilloma accumulation rate of T>A transversions is consistent with previous studies showing that papillomas are polyclonal, whereas SCCs are largely dominated by a single clone ^10^.

These findings demonstrate that longitudinal analysis of hair follicles can provide a unique window into how mutations evolve within the skin cellular architecture.

### Hair follicles record the mutational history of tumour-destined skin regions

After mechanical wounding, hair follicle stem cells migrate to the interfollicular epidermis ^24^. We hypothesised that similar migration would happen after mutagenic exposure. This would predict that hair follicle somatic mutations should be shared with the surrounding epidermis. If so, then our longitudinal approach would reveal the dynamic clonal evolution of every profiled skin region, including those destined to form tumours.

Hair follicles and skin, across the different morphologies, share 2.6% of mutations, while most of the mutations are within the same compartment (Figure 4A; Supplementary Figure 4A, B). Of all mutations identified by exome sequencing in tumour-containing skin biopsies, 12.1% were also detected in hair follicle biopsies collected at one or more earlier timepoints (Figure 4B; Supplementary Table 4). Thus, our hair follicle experiment can be used to trace clonal evolution.

**Figure 4.**
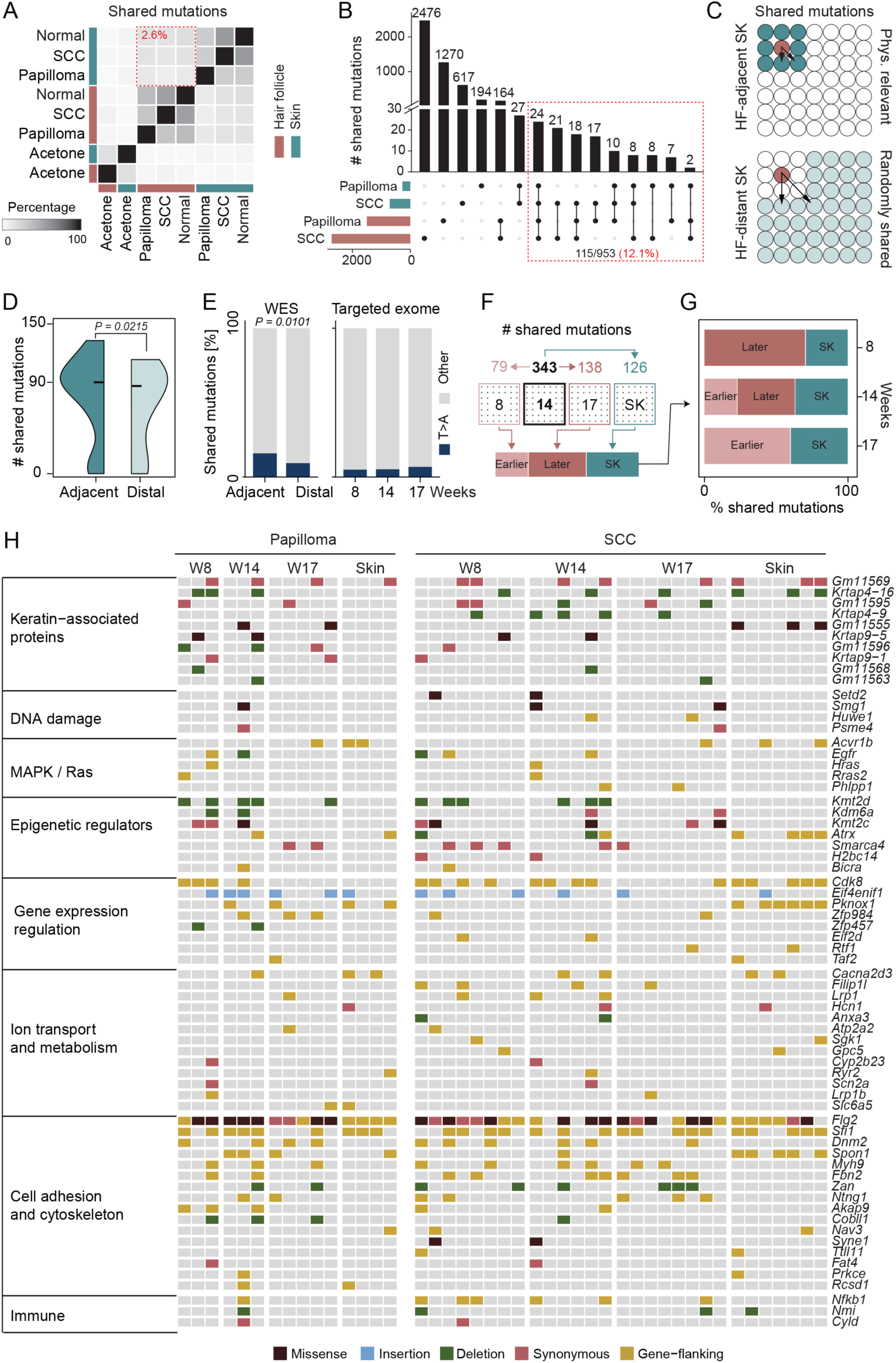
Hair follicles preserve mutational signatures of the epidermis during tumour development. (**A**) Heatmap showing percentage of shared mutations identified in hair follicles (brown) or skin (green) biopsies in normal or tumour-destined (papilloma, SCC) regions obtained from targeted exome sequencing. Hair follicles and skin biopsies treated with acetone serve as a control. Dotted square: Percentage of shared mutations across skin and hair follicles. (**B**) Number of shared mutations identified in hair follicles (brown) or skin (green) biopsies in tumour-destined (papilloma, SCC) regions obtained from targeted exome sequencing. (**C**) Schematic representation of shared mutations between hair follicle (HF) and skin (SK) in spatially adjacent (top) and distal (bottom) regions. Shared mutations detected in adjacent HF-SK pairs are interpreted as potentially physiologically (Phys.) relevant and indicative of shared clonal origin; shared mutations between distal regions are more likely to arise randomly. (**D**) Quantification of shared mutations between HF and SK in adjacent versus distal regions using whole-exome data. P-value: Poisson mixed-effects model controlling for per-biopsy coverage and callable territory (β = 0.12096). (**E**) Proportion of T>A mutations detected by whole exome (WES) in adjacent and distal hair follicle biopsies (left) and targeted exome over time (right). (**F**) Schematic representation of the analysis performed to detect mutations that are shared between earlier and later hair follicle time points and skin. (**G**) Percentage of mutations shared between earlier and later hair follicle time points and skin. (**H**) Oncoprint showing mutated genes shared between hair follicle time points (8, 14, 17 weeks) and skin (19 weeks) in SCC and papilloma-associated regions. Columns correspond to individual biopsies. Mutations are colour-coded by the indicated mutation type.

We reasoned that clonal expansion would be spatially restricted, meaning that adjacent hair follicles and skin should share more mutations than distal regions (Figure 4C). Consistent with this hypothesis, our analysis found more shared mutations between adjacent hair follicles and skin regions than between distal pairs (Figure 4D; Supplementary Figure 4C-D). The mutations shared by widely dispersed regions probably reflect background genetic changes accumulated over time. Importantly, the DMBA-associated T>A transversions were enriched among mutations shared between adjacent regions compared with those shared between distal pairs (Figure 4E).

We traced the fate of shared mutations over time (Figure 4F). Many mutations first detected in hair follicles at early time points persisted throughout disease progression and were subsequently detected in matched epidermal regions (Figure 4G; Supplementary Figure 4E). These persistent mutations mapped to known SCC-associated genes (Figure 4H; Supplementary Figure 4F, G), such as *Hras* and *Egfr*, both established drivers of human SCC progression ^3,25^. The gene set was dominated by chromatin regulators recurrently inactivated in human cutaneous squamous carcinoma (*Kmt2d*, *Kdm6a*, *Smarca4* and *Atrx*) whose loss compromises both homologous recombination and mismatch repair ^26–28^. We also recovered core components of the NF-κB axis (*Cyld* and *Nfkb1*) whose keratinocyte-specific disruption is sufficient to accelerate skin tumorigenesis ^25,29^. A second group of persistently mutated genes, with no established role in squamous carcinogenesis, includes *Psme4*, which encodes a proteasome activator that mediates acetylation-dependent core histone degradation following DNA damage ^30^.

Thus, longitudinal analysis of DNA isolated from hair follicles can provide dynamic insight into the evolutionary history of the underlying skin regions and provide early information on the acquisition and expansion of cancer-related mutations.

### Early hair follicle mutations identify tumour initiation and predisposition candidate genes

Using the longitudinal tracking of shared hair follicle-skin mutations, we then identified candidate genes associated with tumour initiation or predisposition. We classified recurrently mutated genes in both hair follicles and matched skin at the earliest stage they were detected and associated them to tissue morphology. This identified three classes of genes: First, initiation genes (I) were defined as genes harbouring mutations exclusively in tumour-destined samples, with no mutations detected in normal or control samples. Second, predisposition genes (P) were defined as genes harbouring mutations in both non-regressing tumours and macroscopically normal regions across hair follicle and skin samples. Third, genes fulfilling both criteria (I+P) (Figure 5A; Supplementary Figure 5A-C).

**Figure 5.**
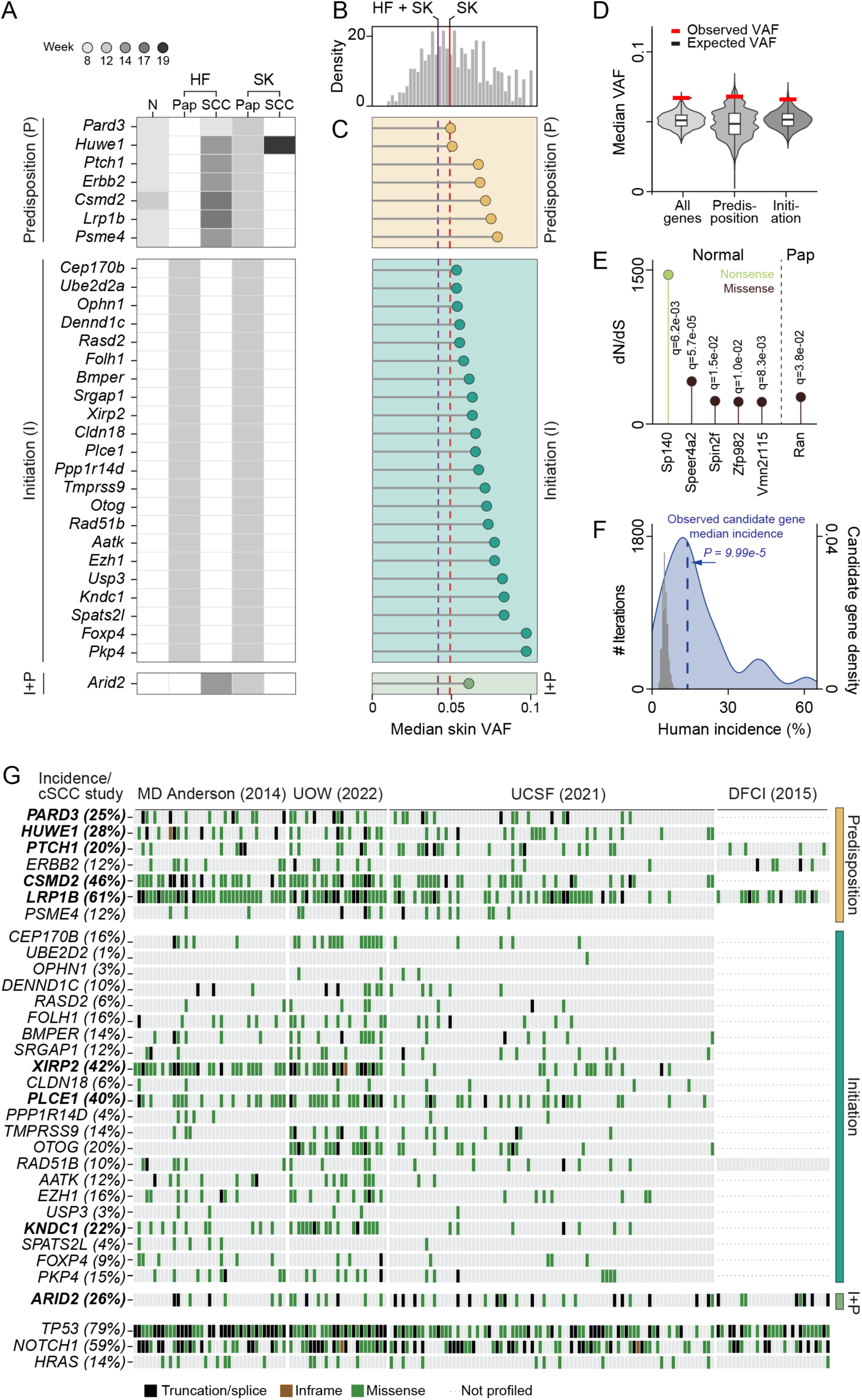
Longitudinal hair follicle profiling identifies candidate genes associated with tumour initiation and predisposition. **(A)** Recurrently mutated genes in macroscopically normal (N) or tumour-destined (SCC, papilloma) regions found in hair follicles (HF) and skin biopsies (SK). Genes are grouped according to their potential relevance for tumour initiation (I), predisposition for malignant transformation (P), or both (I+P). Shaded grey: Time point at which the mutation was first detected. (**B**) Distribution of variant allele frequencies (VAFs) of all mutations identified (grey bars). Red line: Median of VAF distribution mutations found only in skin (SK). Purple line: Median of VAF distribution mutations shared by skin and hair follicles (HF+SK). (**C**) Median VAF of predisposition (orange), initiation (green) and both categories (light green) candidate genes shown in (A). (**D**) Distribution and density of median VAFs from permuted gene sets for each candidate-gene group (all, initiation, and predisposition). Red bar: Observed median VAF from each candidate gene group. Boxplots summarise the median and interquartile range of each distribution. (**E**) Positive selection analysis based on the ratio of nonsynonymous (dN) to synonymous (dS) substitution rates in macroscopically normal skin (Normal) and papillomas (Pap). Green dot: Nonsense mutation rate. Black dot: Missense mutation rate. Q-values (Benjamini-Hochberg adjusted) are shown above each gene. (**F**) Incidence of candidate-gene mutations shown in (G) in human cSCC. Grey bars: Distribution of median mutation incidences of 10,000 size-matched random gene sets. The bar height indicates the number of iterations in each incidence bin. Blue curve: Smoothed density estimate of the same randomized distribution of median incidence of size-matched random gene sets. The dashed blue line: Observed median incidence of the candidate genes (P = 9.99 × 10⁻⁵). (**G)** Somatic mutations in candidate genes across four independent human cSCC cohorts. Columns represent individual tumours and rows represent genes; mutation classes are indicated as truncation/splice (black), in-frame indels (brown), missense (green), or not profiled (_•••_). Percentages indicate the proportion of tumours harbouring a mutation in each gene. Genes are grouped by their proposed stage of involvement, in tumour predisposition and initiation with established cSCC drivers (*TP53, NOTCH1*, and *HRAS*) shown for comparison. Bold gene names denote longitudinally identified genes in hair follicles showing relatively high mutation incidences in human cSCC.

Many genes associated with predisposition have previously been implicated in human skin cancer, including the tumour suppressors *Ptch1 and Pard3* ^31,32^, as well as the RAS-associated oncogene *Huwe1* ^33^. Others are known from different cancers, such as *Csmd2* in gastric cancer ^34^. Similarly, initiation genes include *Rad51b*, a germline susceptibility locus for skin cancer ^35^. Others include the recurrently mutated *Plce1* and *Xirp2* gastric cancer genes ^36,37^, and *Cldn18*, whose fusion with *ARHGAP26* defines a molecular subtype of diffuse gastric carcinoma ^38^. Finally, our identification of *Arid2* as the only gene shared between the predisposition and initiation categories is consistent with its known biology as established tumour suppressor in melanoma ^39^.

To identify whether our candidate genes showed hallmarks of positive selection, we compared their variant allele frequencies (VAF) with randomly sampled gene sets. As a reference, we first determined the VAF distribution of all skin mutations, as well as the median VAF for mutations found only in skin (SK) and those shared by skin and hair follicle(s) (HF+SK) (Figure 5B). All candidate genes exhibited higher median VAF than both median VAF references (Figure 5C; Supplementary Table 5). To test whether the median VAF of our predisposition and initiation genes was higher than background, we determined a background distribution by performing 10, 000 median VAF calculations for randomly chosen sets of genes of the same size (Methods). Indeed, the observed VAFs for the candidate genes were consistently higher than expected (Figure 5D), indicating that clones carrying the candidate genes preferentially expanded during tumorigenesis.

When considering all mutated genes, we identified a correlation, albeit weak, between gene length and absolute number of mutations. However, the initiation and predisposition genes showed a clear enrichment in total mutations above that expected from their length alone (Supplementary Figure 5D). Among the specific types of mutational disruption, missense and splicing were statistically enriched compared to the permuted gene sets (empirical *P* = 0.015 and *P* = 0.007, respectively; Methods; Supplementary Figure 5E, F), demonstrating that longitudinal hair follicle analysis identified genes recurrently altered in papillomas and SCC.

To further characterise genes under positive selection in skin, we measured the ratio of nonsynonymous to synonymous mutations (dN/dS; Figure 5E). We identified five significantly positively selected genes in macroscopically normal tissue (*Sp140, Speer4a2, Spin2f, Zfp982*, and *Vmn2r115*), none of which are known to be involved in skin tumorigenesis. In contrast, papillomas displayed positive selection for a single gene, *Ran*, a regulator of nuclear transport and cell division known to be involved in tumour progression and metastasis ^40^. Finally, we directly searched for evidence of mutations occurring in *Hras, Kras,* and *Nras*, members of the MAPK pathway, which are frequently mutagenized in human SCC ^7^ and in DMBA/TPA-driven SCC ^3^. In all three genes, we found multiple known mutations, including *Hras^Q61L^* (Supplementary Figure 5G) ^3^.

### Longitudinally identified genes are mutated in corresponding human skin cancers

Finally, we compared our candidate genes with recurrently mutated genes found in cSSC patient cohorts by re-analysing datasets from the DFCI ^41^, MD Anderson ^42^, UCSF ^43^, and UOW ^44^ studies. All candidate genes harboured mutations across these independent human datasets, with an average mutation incidence of 14%, representing a 2.7-fold enrichment relative to the expected background rate (5.1%) (Figure 5F, G; Supplementary Figure 5H). Among the frequently mutated genes in the human cohorts with known roles in skin cancer, we identified genes that we had defined as predisposition (*PARD3, PTCH1*) ^32,33,42^ and initiation (*PLCE1, ARID2*) ^39,45^ genes. The initiation gene *KNDC1*, which encodes a Ras guanine nucleotide exchange factor, was also frequently mutates in the human cohorts, yet has not previously been implicated in cSCC. Notably, the previously established cSCC driver gene *HRAS* ^3,6^ showed a similar mutation incidence in mouse and human (Figure 5G; Supplementary Figure 5G).

Importantly, we detected mutations in many of these genes in macroscopically normal tissue, long before the appearance of overt lesions. Thus, longitudinal hair follicle profiling can prospectively identify genomic alterations associated with future tumour development. The recurrence of these alterations across independent human cohorts suggests that the evolutionary trajectories captured by our mouse model converge on genes under positive selection during human squamous cell carcinogenesis. Together, these findings establish longitudinal hair follicle analysis as a non-invasive framework for identifying tumour-prone regions and evolutionarily selected candidate genes, long before overt malignancy.

## Discussion

During cutaneous carcinogenesis, initiated clones persist and expand across visibly normal skin long before a lesion appears. We show that hair follicles can be used as a non-invasive genomic readout for longitudinally tracing clonal evolution and pinpointing tumour-prone fields before malignant transformation.

Hair follicle stem cells serve as the cells of origin for tumour initiation and subsequent clonal expansion in SCC ^46^. Consistent with this, mutations detected in hair follicles recurred in matched papilloma-and squamous cell carcinoma-associated skin, showing that the two compartments share clonal lineages throughout tumorigenesis and that skin lesions can originate in the hair follicle. Crucially, shared mutations were enriched in spatially adjacent rather than distal regions: follicles retain the clonal history of their proximal epidermis, and can therefore delineate the field of cancerization.

Hair follicles carried an elevated mutational burden that declined during tumorigenesis, unexpected for a slow-cycling compartment ^15,20^. After mutagenic injury, bulge stem cells bearing DNA lesions mount an early post-transcriptional response that suppresses proliferation, DNA repair, and cell-cycle machinery, reinforcing quiescence rather than triggering repair-coupled proliferation. Slow cycling is thought to limit mutagenesis by minimising replication error; here, the same quiescence-reinforcing program also restrains repair, allowing damage to persist unresolved. Studies in haematopoietic stem cells show that DNA damage can accumulate during quiescence and then be resolved upon cell-cycle re-entry ^47^. Similarly, in the bulge stem cell lesions are fixed into the genome when the cell re-enters the cycle during hair-cycle progression and TPA-driven proliferation, and thus bulge cells accumulate mutations acting as a reservoir. Meanwhile, fewer mutations in the IFE could be explained by loss of cells due to shedding of the outermost layers ^48^.

Hair follicles in tumour-destined regions harboured the most mutations at the pre-malignant stage of carcinogenesis, suggesting the presence of multiple coexisting heterogeneous clones at the benign stage, as described in skin papillomas ^10^. The subsequent decline reflects clonal selection, as diverse early clones are replaced by a few expanding tumour-associated lineages. This resembles the clonal sweep observed in invasive squamous cell carcinoma, where tumour growth can become dominated by a single clone. The decline in mutational burden is therefore consistent with clonal selection and expansion in both hair follicles and skin.

The spatiotemporal resolution of our study enabled genes to be assigned to carcinogenic stages: mutations shared between normal and later tumour-associated samples mark predisposition events, those shared between papilloma-associated hair follicles and skin mark benign initiation, and those retained from papilloma to SCC mark progression. Our cross-species comparison supports the biological relevance of these clonal trajectories, as candidate genes identified from early, pre-malignant hair follicle mutations recurrently harboured mutations also found across independent human cSCC cohorts. This extends the finding that macroscopically normal human epidermis already harbours cancer-driver clones under strong positive selection at a given moment ^49^. Indeed, our study newly reveals how clones carrying cancer drivers can be tracked prospectively from a non-invasive, repeatedly sampleable tissue, ahead of malignant transformation.

Hair follicles are readily accessible in routine dermatology and can be sampled with minimal invasiveness. Sampling them from individuals at elevated risk could flag tumour-prone regions years before a lesion becomes visible, opening a practical route to non-invasive skin-cancer risk detection and prevention.

## Material and Methods

### Two-stage skin carcinogenesis model

Multi-stage chemical carcinogenesis in mouse skin was conducted at the DKFZ animal facility in accordance with institutional animal care guidelines and under the terms and conditions of the experimental animal project licences G-96/21 following approval by the local governmental ethics committee. Female C57BL/6J mice (Janvier Labs) were housed under a 12 h light / 12 h dark cycle with ad libitum access to water, standard chow, and environmental enrichment.

For the two-stage chemical carcinogenesis protocol ^2^, the dorsal skin of mice was subjected to a single topical application of 100 µg 7,12-dimethylbenz[a]anthracene (DMBA; Sigma-Aldrich) followed by repeated exposure to 10 µg 12-O-tetradecanoylphorbol-13-acetate (TPA; Sigma-Aldrich) three times per week until the experimental endpoint. Animals were euthanised at week 12 or week 19 to collect samples for exome sequencing and at week 10-12 to collect samples for single-cell RNA sequencing.

For short-term mutagen exposure, 7-week-old female C57BL/6J mice received a single topical application of 500 µg DMBA dissolved in 200 µl acetone. Control animals were treated only with acetone. Mice were euthanised 48 h after application.

### Transgenic mice

Rosa-CAG-LSL-tdTomato ^50^, K19-CreER ^51^, Lgr5-CreERT2 ^52^, Lrig1-EGFP-ires-CreERT2 ^53^, Lgr6-EGFP-ires-CreERT2 ^54^ mice have been described previously. Mice were either housed in the Wellcome Trust-Medical Research Council Cambridge Stem Cell Institute Animal Unit or the DKFZ Central Animal Laboratory. Mouse husbandry and experiments were carried out according to the local ethics committee under the terms of a UK Home Office license P36B3A804 approved by the Animal Welfare and Ethical Review Body (AWERB) or in accordance with guidelines of the local ethics committee under the terms and conditions of the animal licences G-41-21 approved by the ‘Regierungspräsidium Karlsruhe’.

To conditionally induce tdTomato-reporter lines for expression of Cre recombinase, Rosa-CAG-LSL-tdTomato mice were crossed with K19-CreER, Lgr5-CreERT2, Lrig1-CreERT2 and Lgr6-CreERT2 mice. To activate CreER, mice were treated 7 days before UVB irradiation with a single intraperitoneal (i.p.) injection of 100 ug (Lrig1), 500 ug (Lgr5, K19) or 1000 ug (Lgr6) of tamoxifen (Sigma-Aldrich) diluted in corn oil. UVB exposure was performed on the dorsal skin of 8-week-old mice. Mice were treated with 2.5 kJ/m^2^ UVB (310 nm) irradiation.

## Longitudinal hair sampling

To enable spatially matched longitudinal sampling, a 25 × 25 mm grid was tattooed onto the dorsal skin of each mouse using needle-delivered ink. At each time point, the grid area was stripped using duct tape. A sampling grid was then marked on the tape, and the marked regions were punched (3 mm) for hair follicle collection. Hair follicles were collected at weeks 8, 14, 17 and 19 for targeted exome sequencing, and at week 12 for whole-exome sequencing. At the end of the experiment (week 19), the corresponding dorsal skin area was collected post-mortem using 3-mm punch biopsies.

Hair follicle and skin samples were annotated based on the macroscopic lesion status of the corresponding grid position. Regions that subsequently developed squamous cell carcinoma were retrospectively classified as ‘SCC’ from the earliest collection timepoint, including samples obtained before carcinoma formation. Regions associated with papillomas or regressing non-carcinoma lesions were classified as ‘Papilloma’, whereas regions without visible lesions throughout the experiment were classified as ‘macroscopically normal’.

### DNA extraction

The 3 mm tape punches were transferred into 96-well plates containing Qiagen ATL buffer and proteinase K to proceed with DNA extraction. At the experimental endpoint, the skin containing the tattooed grid was excised and transferred onto a polystyrene platform to preserve spatial orientation. Matching 3 mm skin punches were collected and transferred into 96-well plates. Samples were snap-frozen in liquid nitrogen and stored at -80°C until DNA extraction. At the experimental endpoint, the liver was also collected, minced into small pieces, snap-frozen in liquid nitrogen and stored at -80°C until DNA extraction.

Hair follicle DNA was extracted using the Qiagen DNeasy workflow. Hair follicles in the 96-well plates were released from the tape by mechanical scraping with the flat plastic end of a syringe and incubated overnight at 56°C. Following digestion, samples were briefly centrifuged to collect the condensate, supplemented with 200 µl of Buffer AL, mixed thoroughly, and incubated at 56°C for 30 min. Subsequently, 200 µl of ethanol was added, and lysates were transferred to DNeasy spin columns. Columns were centrifuged, sequentially washed with 500 µl of AW1 and 500 µl of AW2 buffers, and dried by an additional centrifugation step to remove residual ethanol. DNA was eluted in 25 µl pre-warmed AE buffer; the eluate was re-applied to the column for a second elution to maximise DNA recovery.

DNA from skin biopsies and livers was extracted using a combination of the AllPrep DNA/RNA Micro Kit (Qiagen) and DNeasy Blood and Tissue Kit (Qiagen). Briefly, 350 μl of RLT buffer (from the AllPrep DNA/RNA Micro Kit) containing 1% β-mercaptoethanol (Sigma-Aldrich) was added to each sample, and the tissue was homogenised in a Qiagen TissueLyser II for 2 cycles of 3 minutes at 30 Hz. After lysis, nuclease-free water and proteinase K (from DNeasy Blood and Tissue Kit) were added to the lysate and incubated at 56°C for 10 minutes. Buffer AL (Qiagen Blood & Tissue) was added, mixed by vortexing, and incubated at 56°C for 30 minutes at 500 rpm. An equal volume of absolute ethanol was added to the samples and mixed thoroughly. The precipitated mixture was transferred to a DNeasy Mini 96-well spin plate (Qiagen), washed twice with 700 μl buffer AW1 and twice with buffer AW2. After drying the membrane for 3 minutes at 20,000 x g, DNA was eluted using two consecutive elutions of 30 µl pre-warmed AE buffer (70°C). DNA yield and fragment distribution were assessed using Qubit and TapeStation HS dsDNA assays.

### Targeted exome sequencing

Targeted exome sequencing libraries were generated using the Agilent SureSelect XT Low Input enzymatic fragmentation workflow with dual indexing. Briefly, 10-200 ng genomic DNA was used as input where available. DNA was enzymatically fragmented, with fragmentation times adjusted according to tissue type: 20 min for skin-derived DNA and 27 min for hair follicle-derived DNA. Fragmented DNA was subjected to end repair and dA-tailing, followed by ligation of P5-indexed adapters. Adapter-ligated libraries were purified using AMPure XP beads and amplified by pre-capture PCR. Pre-capture library quality and size distribution were assessed using Agilent TapeStation D1000 ScreenTape, and library concentration was calculated by integrating the 200-400 bp fragment range.

For target enrichment, purified pre-capture libraries were hybridised to the custom-designed capture probe library using the SureSelect hybridisation workflow. The custom-designed panel contained 225 genes and was 1.9 Mb in size, without UTR and with 50 bases on either side of the exons (design ID: 3396321). Hybridised DNA was captured using Dynabeads MyOne Streptavidin T1 magnetic beads, followed by stringent washing with SureSelect wash buffers. Captured libraries were then amplified and purified using AMPure XP beads. Final libraries were quantified and quality-controlled using High Sensitivity D1000 ScreenTape, and samples were pooled in equimolar amounts for multiplexed sequencing.

### Whole-exome sequencing

Dual indexed whole exome capture libraries were prepared using SureSelect XT Low Input Enzymatic Fragmentation protocol, as described above (Agilent). For mouse whole exome sequencing (WES) libraries SureSelectXT Low Input Target Enrichment System was used with Mouse All Exon capture kit (52 Mb - cat. No. 5190-4642). Exome libraries were sequenced on Illumina NovaSeq sequencers using a 100 bp paired-end read protocol.

### Single-cell isolation

Shaved dorsal skin was collected as 1-5 mm³ biopsies and dissociated using one of two sequential enzymatic digestion procedures. In the first procedure, tissue samples were incubated for 2.5 h in HBSS (Invitrogen) containing 1.25 mg ml⁻¹ collagenase I (Sigma), 0.5 mg ml⁻¹ collagenase II (Worthington), 0.5 mg ml⁻¹ collagenase IV (Worthington), and 0.1 mg ml⁻¹ hyaluronic acid (Sigma). Trypsin (Santa Cruz Biotechnology) was then added to a final concentration of 0.25%, and incubation was continued for an additional 30 min. Alternatively, biopsies were first treated with 0.25% trypsin for 2 h and subsequently incubated for 1 h in the collagenase-containing dissociation solution described above. All enzymatic digestion steps were carried out at 37 °C with agitation at 600 rpm.

Following a total digestion time of 3 h, enzyme activity was stopped by adding HBSS supplemented with 10% fetal bovine serum (FBS; Life Technologies). The resulting cell suspensions were passed through 40 µm cell strainers (Greiner Bio-One), which were subsequently rinsed with an additional 2–3 ml of HBSS containing 10% FBS. Cells were collected by centrifugation at 300 × g for 8 min at 4 °C. Pellets were gently resuspended using wide-bore P1000 pipette tips in 100–200 µl of sorting buffer composed of PBS containing 1 mM EDTA (Invitrogen) and 0.04% HyClone bovine serum albumin (BSA; GE Healthcare Life Sciences). Suspensions were either filtered once more through 40 µm FLOWMI cell strainers (Fisher Scientific), followed by assessment of cell concentration and viability using 0.4% trypan blue (Invitrogen) and a Countess II automated cell counter (Invitrogen), or processed by fluorescence-activated cell sorting (FACS).

For FACS, cells were resuspended in sorting buffer supplemented with 1 µg ml⁻¹ propidium iodide (PI; Life Technologies) and transferred to low-binding polypropylene tubes. Viable cells from each sample were sorted with a BD Aria II cell sorter (BD Biosciences) equipped with a 70 µm nozzle operated at 70 psi. Cells were collected into individual wells of a flat-bottom, low-attachment 96-well tissue-culture plate containing 30 µl of sorting buffer and maintained on a chilled plate. PI was excited using a 561 nm laser, and emitted fluorescence was detected through a 610/20 nm bandpass filter preceded by a 595 nm long-pass filter. Debris was removed by gating on forward-scatter area versus side-scatter area, doublets were excluded using side-scatter width versus side-scatter area, and autofluorescent or PI-positive cells were excluded based on RL780/60-A versus PI-A. Samples were maintained on ice throughout processing.

### Single-cell RNA-sequencing

Single-cell 3′ gene-expression libraries were generated using the 10x Genomics Chromium platform in accordance with the manufacturer’s protocol. For each 1 mm³ skin biopsy, the complete sorted-cell suspension was loaded onto the Chromium Controller, corresponding to approximately 800-1,100 cells from untreated samples and 2,900-5,300 cells from DMBA/TPA-treated samples. Cell suspensions obtained from 5 mm³ biopsies were diluted to achieve an anticipated recovery of approximately 5,000 cells.

Individual cells were partitioned into droplets using the Chromium Controller, in which cell lysis and barcoded reverse transcription of mRNA were performed. The resulting cDNA was subsequently amplified and fragmented, followed by construction of sequencing libraries compatible with Illumina platforms. Library concentrations were measured using the Qubit dsDNA High Sensitivity Assay Kit (Life Technologies), while cDNA quality and fragment-size distributions were evaluated using D1000 ScreenTapes (Agilent Technologies).

Libraries were pooled and sequenced using an Illumina NextSeq 2000, NovaSeq 6000, or NovaSeq X Plus instrument. Paired-end sequencing was performed using either a 28-bp read 1 and a 94-bp read 2 configuration or 100-bp reads for both read 1 and read 2.

### RNA extraction of UVB exposed epidermal compartments

Mouse dorsal skin was isolated as previously described ^55^. To isolate specific epidermal cell populations, dissociated epidermal cells were stained with DAPI, PE-conjugated anti-ITGA6, FITC-conjugated anti-CD34, PE-Cy7-conjugated anti-CD31, PE-Cy7-conjugated anti-CD117, and PE-Cy7-conjugated anti-CD45 antibodies, followed by fluorescence-activated cell sorting (FACS). DAPI-positive and PE-Cy7-positive cells were excluded. Cells of double positive staining for PE-ITGA6 and FITC-CD34 were collected as hair follicle bulge cells, whereas PE-ITGA6 positive only cells were collected as basal layer interfollicular epidermal cells.

Total RNA was extracted using the Direct-zol RNA Microprep Kit (Zymo Research, R2062) according to the manufacturer’s instructions. RNA-seq libraries were constructed using the Illumina Stranded Total RNA Prep, Ligation with Ribo-Zero Plus kit (Illumina, #20040529) according to the manufacturer’s instructions. The prepared RNA-seq libraries were sequenced on an Illumina NovaSeq 6000 system using an S4 flow cell.

### Histology, immunostaining, antibodies and imaging

Tissues were either embedded in OCT and frozen, or fixed overnight in 4% paraformaldehyde, transferred to 70% ethanol and embedded in paraffin. Sections were cut at 4–5 µm (paraffin) or 10–50 µm (frozen). Immunofluorescence and immunohistochemistry staining of frozen and paraffin-embedded tissues were performed essentially as described previously ^56,57^. Sections were deparaffinized in xylene and rehydrated through 100%, 70% and 40% ethanol into distilled water. Antigen retrieval was performed at 98 °C for 30 min in pH 9.0 Tris-EDTA buffer (Abcam, ab93684) supplemented with 0.05% Tween-20. After cooling for 40 min, sections were blocked for 1 h at room temperature in 10% FBS, 0.05% Tween-20. Primary antibodies were diluted in blocking buffer and incubated overnight at 4 °C. Sections were then washed three times for 5 min in PBS and incubated for 1 h at room temperature in the dark with secondary antibodies and DAPI diluted in blocking buffer. Sections were washed and mounted either in DAKO mounting medium (Agilent) or in glycerol supplemented with Mowiol 4-88 (Sigma-Aldrich).

Primary antibodies: anti-Ki67 (1:200; clone SP6, Vector Laboratories, VP-RM04); anti-Ki67 (1:50; Abcam, ab16667); anti-p53 (1:200; Vector Laboratories, VP-P956); anti-CPD (1:2000; CosmoBio, NMDND001); rat monoclonal anti-CD34 (1:50; eBioscience, 14-0341-81); rabbit monoclonal anti-phospho-histone H2A.X (1:50; Cell Signaling Technology, 9718S). Secondary antibodies: goat anti-rabbit Alexa Fluor Plus 647 (1:1000; Invitrogen, A32733); goat anti-rat Alexa Fluor 488 (1:1000; Invitrogen, A-11006). Nuclei were labelled with DAPI (1:1000; Roche, 10236276001).

Brightfield and fluorescence images were acquired on an Axio Imager 2 upright microscope (Zeiss), a Leica TCS SP8 confocal microscope, or a Zeiss LSM700 confocal microscope. LSM700 acquisition was performed in Z-stack mode (three images over a 4 µm range), and maximum-intensity projections were used for quantification and visualisation. Explant images were acquired on a Leica DMI4000 microscope. Images were processed in Fiji ^58^.

### TUNEL assay

DNA fragmentation associated with apoptosis was assessed in formalin-fixed, paraffin-embedded skin sections using the TUNEL Assay Kit (ab66110; Abcam), with minor modifications to the manufacturer’s protocol. Sections were deparaffinized by two 5-min incubations in xylene and subsequently rehydrated through a graded ethanol series, followed by immersion in 0.85% NaCl and washing in PBS. Tissue sections were permeabilised with Proteinase K for 10 min. For positive controls, sections were additionally treated with DNase I (Qiagen, cat. no. 79254) for 15 min at 37 °C. Following washing in PBS, sections were post-fixed in 4% paraformaldehyde for 5 min and equilibrated in PBS. DNA strand breaks were labelled by incubating the sections with a DNA-labelling mixture containing TdT reaction buffer, TdT enzyme, and Br-dUTP for 1.5 h at 37 °C in a humidified chamber. After washing with PBS, sections were incubated with an anti-BrdU-Red antibody for 30 min at room temperature while protected from light. Negative-control sections were processed identically, except that TdT enzyme was omitted from the labelling reaction. Nuclei were counterstained with DAPI diluted 1:3,000, rinsed with ddH2O, and mounted using an aqueous fluorescence mounting medium (Agilent Technologies).

### RNA sequencing analysis

RNA-sequencing data were processed using the DKFZ One Touch Pipeline (OTP) ^59^ with RNA-seq Workflow version 1.3.0 in combination with the Roddy workflow management system version 3.5.9. Briefly, sequencing reads were aligned to the GRCm38mm10_PhiX reference genome, consisting of GRCm38 supplemented with PhiX contigs, using STAR aligner version 2.5.3a ^60^. PCR/optical duplicates were marked using Sambamba version 0.6.5. Quality control was performed using flagstat from SAMtools version 1.6 and RNA-SeQC version 1.1.8. Gene-level read quantification was performed using featureCounts from the Subread package version 1.5.1 with the GENCODE mouse gene annotation version M12 and strand-specific counting.

### Exome-sequencing analysis

For the alignment and quality control paired-end FASTQ files were aligned to the GRCm38 (mm10) genome with BWA-MEM (V0.7.15) and sorted with SAMtools (V0.1.19). Duplicates were marked with Picard (V1.125), and the reads were indexed using SAMtools. Coverage was calculated with DKFZ’s ODCF in-house coverageQC and QC matrices were generated using FastQC (V0.11.3) and SAMtools flagstat. All steps were carried out with DKFZ’s OTP AlignmentAndQCWorkflows (<u>github.com/DKFZ-ODCF/AlignmentAndQCWorkflows</u>).

Somatic mutations were identified and filtered using GATK Mutect2 and FilterMutectCalls in tumour-normal mode (v4.6.2.0), with liver samples serving as matched controls to remove germline mutations, with default parameters. Mutation annotation was performed using Ensembl Variant Effect Predictor (VEP v115). Only mutations that passed all filters (PASS flag) were used. Variants with coverage ≤ 10 or ≤ 8 from targeted or whole exome sequencing, respectively, were discarded from subsequent analysis. Mutations called in non-chromosomal contigs, the Y chromosome, and mitochondrial variants were filtered out. Variants with very low allelic fraction (VAF ≤ 0.01) were discarded.

The mutational burden was calculated as the number of mutations per callable megabase (Mb). Callable bases were determined using SAMtools (V1.20). A base was considered callable if it had a minimum base quality (BQ) of 20 and a minimum sequencing depth of 10, with reads required to have a minimum mapping quality (MAPQ) of 20.

For statistical comparisons of morphology groups, mutation counts were modelled using negative binomial generalised linear models (glmmTMB v1.1.14). The log-transformed number of callable megabases was included as an offset to account for differences in callable regions between samples. The differences were evaluated using estimated marginal means and trends (emmeans V2.0.0), with multiple-testing correction using the Benjamini-Hochberg method (BH).

For visualisation and hypermutator detection, mutational burden was log_10_-transformed after adding a pseudocount of 0.05. Samples were classified as hypermutators and excluded from downstream analyses if their transformed mutational burden exceeded the upper Tukey fence, defined as Q3 + 1.5 × IQR.

Mutational signatures were profiled using 6 single-base substitution classes rather than the full trinucleotide context due to limited sample size. Mutations were represented in pyrimidine context by converting purine variants to their pyrimidine counterparts. Temporal changes in T>A mutation proportions were analysed using beta-binomial generalised linear models, modelling T>A mutations against all other mutation counts, with log-transformed callable territory included as a covariate. Category-specific trends and pairwise comparisons at weeks 8, 14, and 17 were assessed using estimated marginal means and trends, with a Benjamini-Hochberg correction for multiple testing.

Variant allele frequency (VAF) was calculated separately for samples with at least two detected mutations. For each test group, sample-level density curves were averaged across the VAF range using an unweighted mean, giving each biological replicate equal weight. The mode was defined as the VAF value corresponding to the maximum of the resulting group-level mean density curve.

For the shared mutation analysis, binary presence-absence matrices were constructed at the level of unique single-base mutations (SBS) and mutated genes, stratified by hair follicles, skin and treatment groups. For the SBS analysis, shared mutations within each mouse were defined as variants with identical chromosomal position and base substitution.

For the gene-level analysis, genes were considered mutated if they contained at least one nonsynonymous mutation regardless of the exact chromosomal position, from any individual mouse. Shared and group-specific mutation sets were summarised as intersection sizes and visualised using UpSetR (V1.4.0).

To test whether transformed hair follicles clonally expanded into neighbouring skin regions, we compared mutation sharing between each hair follicle sample and adjacent or distal skin samples using an empirical distal-null model. The hypothesise was that transformed hair follicles would undergo local expansion into no more than one neighbouring skin grid region. Therefore, for each hair follicle, the observed statistic was defined as the maximum number of mutations shared with any adjacent skin sample. Because the number of adjacent and distal skin samples differed between hair follicles, the distal null was generated by sampling distal skin samples with replacement. For each permutation, the number of distal and adjacent HF-skin pairs was matched within each hair follicle, the maximum number of shared mutations was recorded, and these maxima were summed across hair follicles. A one-sided empirical P-value was calculated as the proportion of permutations with a summed maximum shared mutation count equal to or greater than the observed value. As a complementary model-based analysis, a Poisson generalised linear mixed model was fitted to compare shared mutation counts between adjacent and distal HF–skin pairs, using the maximal number of shared mutations per hair follicle in each group. A random intercept per hair follicle sample, to account for repeated comparisons from the same follicle, was included. Callable sequence size and sequencing coverage in the paired HF and skin samples were included as offsets to account for differences in the opportunity for mutation detection.

For tumour initiation and predisposition gene identification, candidate genes with potential roles in tumorigenesis were identified based on the morphological regions in which nonsynonymous mutations were detected. Genes with potential relevance for tumour initiation and clonal transition from the hair follicle to the skin (initiation genes) were defined as genes harbouring at least one nonsynonymous mutation in tumour-associated hair follicle and skin samples, while lacking nonsynonymous mutations in normal or control samples. Genes potentially predisposing clones to progression from a benign state to SCC (predisposition) were identified as genes harbouring nonsynonymous mutations in non-regressing tumour regions of both hair follicle and skin samples. Only genes with skin VAF above the median VAF of all nonsynonymous skin mutations were retained as genes of interest.

To test whether candidate genes exhibited higher clonal representation than expected by chance, we performed a permutation test. For each permutation, genes were randomly sampled without replacement from the universe of genes mutated in skin samples, while matching the recurrence of the candidate genes. For each of 10,000 permutations, the median VAF was calculated. The observed median VAF of the candidate genes was compared with the permutation-derived null distribution, using a one-sided empirical P-value.

To identify genes under positive selection, the nonsynonymous-to-synonymous mutation ratio (dN/dS) was assessed. Since hair follicles were collected in mini-bulks and are collectively non-clonal, the dNdS approach was performed only on skin samples. dNdScv (v.0.0.1.0) was used to identify genes under selection. Briefly, dNdScv models mutation counts per gene and substitution type using a maximum-likelihood framework that corrects for trinucleotide mutational context, gene-specific mutation rates, and variation in sequence composition and the set of genes included in the analysis. Statistical significance was assessed using likelihood ratio tests to identify genes under positive or negative selection, with multiple-testing correction across genes using the Benjamini-Hochberg false discovery rate (FDR).

### Single-cell RNA-sequencing analysis

Sequencing reads were processed using Cell Ranger v.10.0.0 (10x Genomics) and aligned to the GRCm39 (mm39) mouse reference genome. Quality control was performed separately for each biopsy in Seurat v.5.5.0, using a combination of adaptive filtering based on a median absolute deviation (MAD) strategy and fixed cutoffs. Cells were retained if they had log_10_-transformed total mRNA count and log₁₀-transformed detected genes above the lower adaptive cutoffs (median − 2.5 × MAD) while also meeting the fixed minima of at least 300 detected genes and 500 total mRNA molecules. Mitochondrial read fraction also had to be below either the upper adaptive cutoff (median + 2.5 × MAD) or 20%, whichever was smaller. Putative doublets detected by scDblFinder v.1.23.4 were removed.

Gene-expression data were normalised using SCTransform, and PCA was performed on the scaled data. Data from the individual biopsies were then integrated in Seurat using a reference-based reciprocal PCA approach on the 3,000 most variable features, excluding mitochondrial genes. Reference biopsies were selected as those with the highest number of cells for each treatment group. Louvain clustering was performed at a resolution of 0.8.

For a detailed characterisation of the epidermal compartments, epithelial cells were extracted from the complete dataset. Epithelial-cell expression data was subjected to SCTransform, without regression of additional covariates. Principal component analysis (PCA) was performed, and the first 30 principal components were used for sample (re)integration using reciprocal PCA (RPCA). A shared nearest-neighbour graph was constructed using the first 30 dimensions from integrated RPCA, followed by Louvain clustering at a resolution 0.3, and visualization with Uniform Manifold Approximation and Projection (UMAP). FindAllMarkers function was used to identify marker genes in each cluster. Some clusters were manually removed after verification that they contain a low number of counts/genes in comparison to other clusters, contain gene signatures of another non-epithelial cell type and/or have no presence of clear subtype-defining marker genes along with a high contribution of mitochondrial and/or housekeeping genes. The resulting dataset was reclustered using the above-mentioned steps at resolution 0.2. Cluster annotation of epithelial subtype identities was performed manually using known marker genes. To assign hair follicle subtypes more granularly, hair follicle clusters were subset separately and reclustered at resolution 0.3. In the case of “Bulge_ORS”, where the marker genes of both cell types did not support a definite classification, a “transitional” subtype was assigned. Cell type markers used for visualisation of hair follicle cell types with module scores (AddModuleScore function) were defined according to markers as previously described ^61^.

The following markers were used for bulge module score: *Cd34, Postn, Ptn, Calml3, Tgm5, S100a6, S100a4, Wfdc3, Shisa2, Aqp3*; for ORS module score: *Myof, Dclk1, Sbsn, Tmprss7, Sostdc1, Barx2, Pthlh, Gas1, Krt14, BC016579, Sfrp1, Col16a1, Id4, Gnmt, Scrg1, Mfap2, Il11ra1, Tbx1, Angptl7, Fndc1, Tagln3, Edn2, Cxcl12, Acta2, Myl9, Slc29a1*; for Anagen HF module score: *Krt71, Ctsc, Krt28, Mgst1, Krt25, Paqr5, Top2a, Taf13, Gata3, Krt73, Prss53, Slc39a8, Crym, Fbp1, Nkd2, Acer1, Selenbp1, Krt35, Ube2c, Cdca8, Rnaset2a, Cdc20, Rnaset2b, Ccnb1, Birc5, Dapl1, Alox12, Mt4, Gadd45g, Gjb4, Cited4, Mylk3, Rexo2, Sat1, Ptn, Id1, Nfkbia, Krt36, Tagln3, Ly6g6d, Hephl1, Gprc5d, Krtap11-1, Krt31, Krt33a, Plekha4, Rgcc, Foxq1, Odc1, Prr5l, S100a10, Tmsb10, Krt23, Csta1, Fabp4, Aldh1a3, Krt75, Gpnmb, Sct, Sptssb, Padi3, Tnni1, Krtap8-1, Kitl, Krt26, S100a3, Duoxa1, Mt2, Id3, Ier2, Mt1, Ptma, Junb, Tubb5, Ccnd2, Stmn1, Dut, Hmgb1, Cldn10, Dcn, Ran, Rrm2*; Bulge_ORS was expressing a mixture of bulge and ORS markers, therefore was named as a mix (or transitional phase) of these two populations.

### Single-cell differential expression analysis

For differential expression analysis of the single-cell RNA-sequencing data, raw counts were aggregated across cells belonging to the same individual and cell subtype, generating pseudobulk expression profiles. Pairwise comparisons between the relevant experimental groups were performed using DESeq2 v1.46.0 ^62^ with default settings and treatment as a design factor. P values were corrected for multiple testing using the Benjamini–Hochberg method.

### Gene set enrichment analysis

Genes were ranked by log_2_FC and gene set enrichment analysis was performed using fgsea version 1.32.4 ^63^, restricting gene set size to 10-500 genes. Hallmark, Reactome and GOBP collections were obtained from the mouse MsigDB database ^64–66^ using msigdbr package (https://rdocumentation.org/packages/msigdbr/versions/26.1.0), whereas KEGG LEGACY gene sets were obtained from the human MsigDB collection. Within each cell type and treatment comparison, P values were adjusted separately for each gene-set collection using the Benjamini-Hochberg method. Gene ontology analysis on UVB treated samples was performed using eVITTA v1.3.1. ^67^

### Cell type proportions

To visualize cell type composition across treatments, we analysed epithelial and hair follicle subtypes originating from tissue samples for which trypsin was used as the first dissociation enzyme, which optimizes epithelial cell recovery. Cell type proportions were calculated per biopsy, with each subtype expressed as a percentage of the total analyzed cells within the corresponding epithelial or hair follicle subset. Total numbers of analyzed epithelial or hair follicle cells per biopsy are shown separately on a log10 scale. Cell type proportions were calculated per biopsy, with each subtype expressed as a percentage of the total analyzed cells within the corresponding epithelial or hair follicle subset. Total numbers of analyzed epithelial or hair follicle cells per biopsy are shown separately on a log_10_ scale.

## Mutated human gene analysis

Mouse gene symbols were mapped to human orthologs using the babelgene R package (V22.9) with mouse specified as the source species. Orthologue mapping was performed in two tiers: genes were first queried using a high-confidence threshold (min_support = 3), and genes without a mapping at this level were subsequently queried using a relaxed threshold (min_support = 1). Where multiple orthologs were identified, the top match was retained.

The human orthologs of the 30 candidate genes were evaluated for mutation incidence in human cutaneous squamous cell carcinoma using cBioPortal. Mutation data were compiled from four cSCC studies encompassing 176 patient tumour data (DFCI: cscc_dfarber_2015, MD Anderson: cscc_hgsc_bcm_2014, UCSF: cscc_ucsf_2021, and UOW: cscc_ranson_2022).

For each gene, mutation incidence was calculated as the percentage of profiled tumours harbouring at least one mutation in the corresponding human orthologue, using the number of tumours in which that gene was profiled as the denominator. To test whether candidate genes exhibited higher mutation incidence than expected by chance, a permutation test with 10,000 iterations was performed. For each permutation, 30 genes were sampled without replacement from the background gene set (18,517 mutated genes from all cohorts), and the mean mutation incidence was calculated. The observed median incidence of the candidate genes was compared with the permutation-derived null distribution using a one-sided empirical P-value.

## Acknowledgements

We thank all members of the Odom and Frye lab; the DKFZ Central Animal Laboratory and Next Generation Sequencing core facilities. We are thankful to the Center for Model System and Comparative Pathology (CMCP), Institute of Pathology, University Hospital Heidelberg and the Preclinical and Translational Core Facility, DKFZ, for their services and support in tissue dehydration and sectioning the paraffin-embedded tissues. This research was financially supported by core funding from the Helmholtz Association, the Deutsches Krebsforschungszentrum (DKFZ) and a grant from the European Research Council (788937 to D.T.O.). This work was funded by grants to MF by the Helmholtz Association (W2/W3-106) and Cancer Research UK (CR-UK; C10701/A15181).

## Author Contributions

SDP, AG, MF conceived the study and designed experiments. SDP performed mouse experiments and managed the mouse colonies. SDP performed experiments with assistance from MBeh, MBek, MLK, FC. FX, MP performed the tracing and UV exposure experiments. SDP and MF analysed data. MBeh performed single-cell DMBA experiment with assistance from MBek, MLK, SDP. SDP, YAG, SW, MBeh, MBek designed and implemented computational analysis with assistance from RC and AH-M. SDP, YAG, MF generated figures. MF, DTO supervised the project. SDP and MF wrote the manuscript. All authors had the opportunity to edit the manuscript and approve the final manuscript.

## Data Availability

All single-cell RNA-sequencing data generated in this study have been publicly deposited. The 11-week single-cell RNA-sequencing datasets from DMBA/TPA-treated and acetone-treated (control) mouse back skin are available through the European Nucleotide Archive (ENA) under accession numbers PRJEB58291 and PRJEB58292. The 48-hour single-cell RNA-sequencing dataset from DMBA-treated and acetone-treated mouse back skin is available through ArrayExpress under accession number E-MTAB-16496. The targeted and whole-exome sequencing data are available through ArrayExpress under accession number E-MTAB-17446. The bulk RNA sequencing data after UVB treatment are available through ArrayExpress under accession number E-MTAB-17524.

---

**Supplementary Figure 1.**
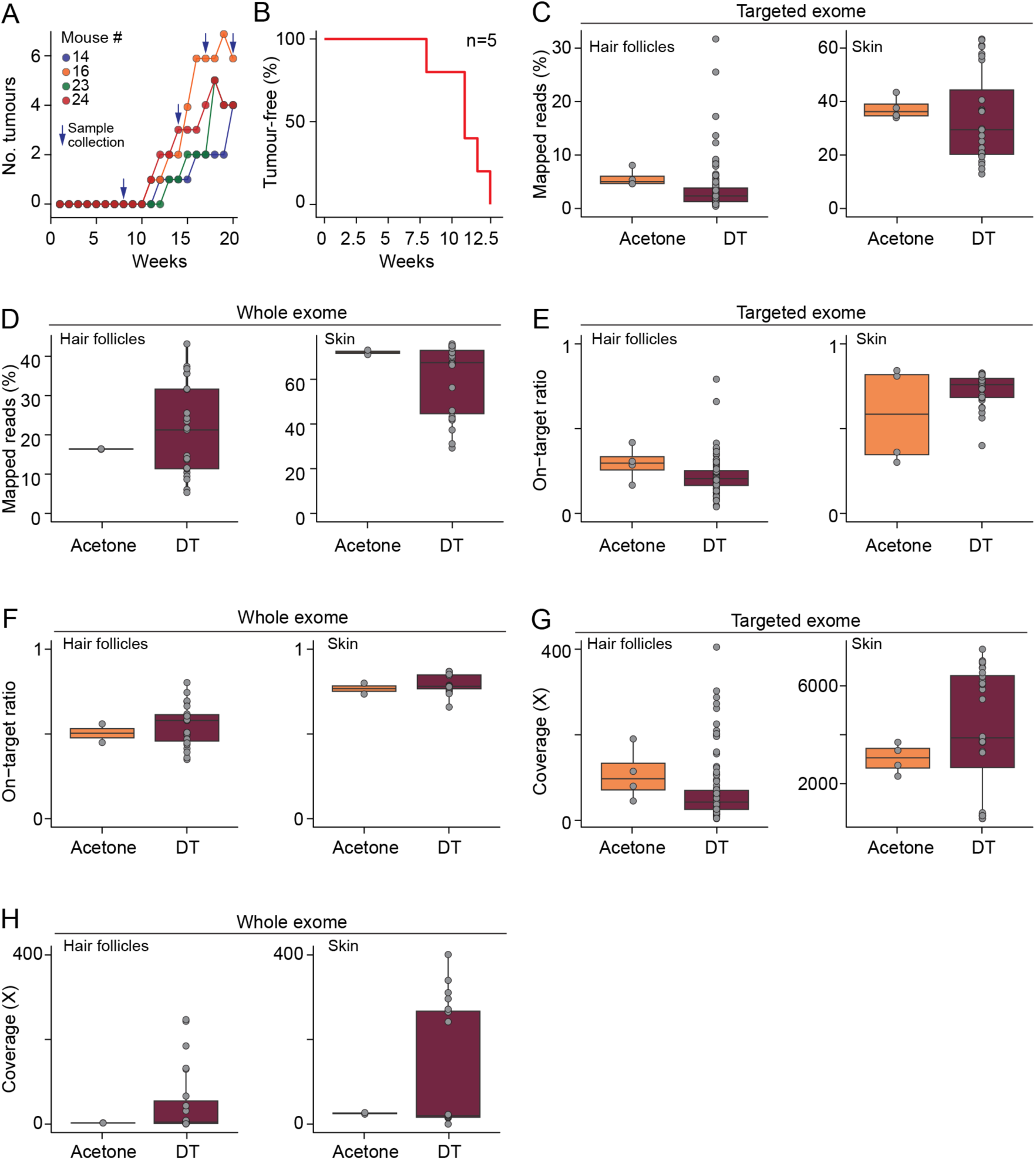
Targeted and whole exome sequencing of hair follicles captured by repeated tape stripping. (**A**) Quantification of tumour formation per mouse over time during DMBA/TPA treatment. (**B**) Kaplan-Meier curve showing the proportion of tumour-free mice over time. n= 5 mice. (**C, D**) Percentage of mapped reads following targeted (C) and whole-exome (D) sequencing. (**E, F**) On-target ratio, defined as the percentage of sequencing reads mapping to the targeted panel (225 genes; 2 Mb) (E) and to the whole exome (F). (**G, H**) Mean number of unique sequencing reads (x) aligning to a given base pair within the target region (coverage). All plots were obtained from hair follicle and skin biopsies from control (acetone) and treated (DT) groups. Box plots indicate the median and interquartile range; each point represents an individual biopsy grid sample.

**Supplementary Figure 2.**
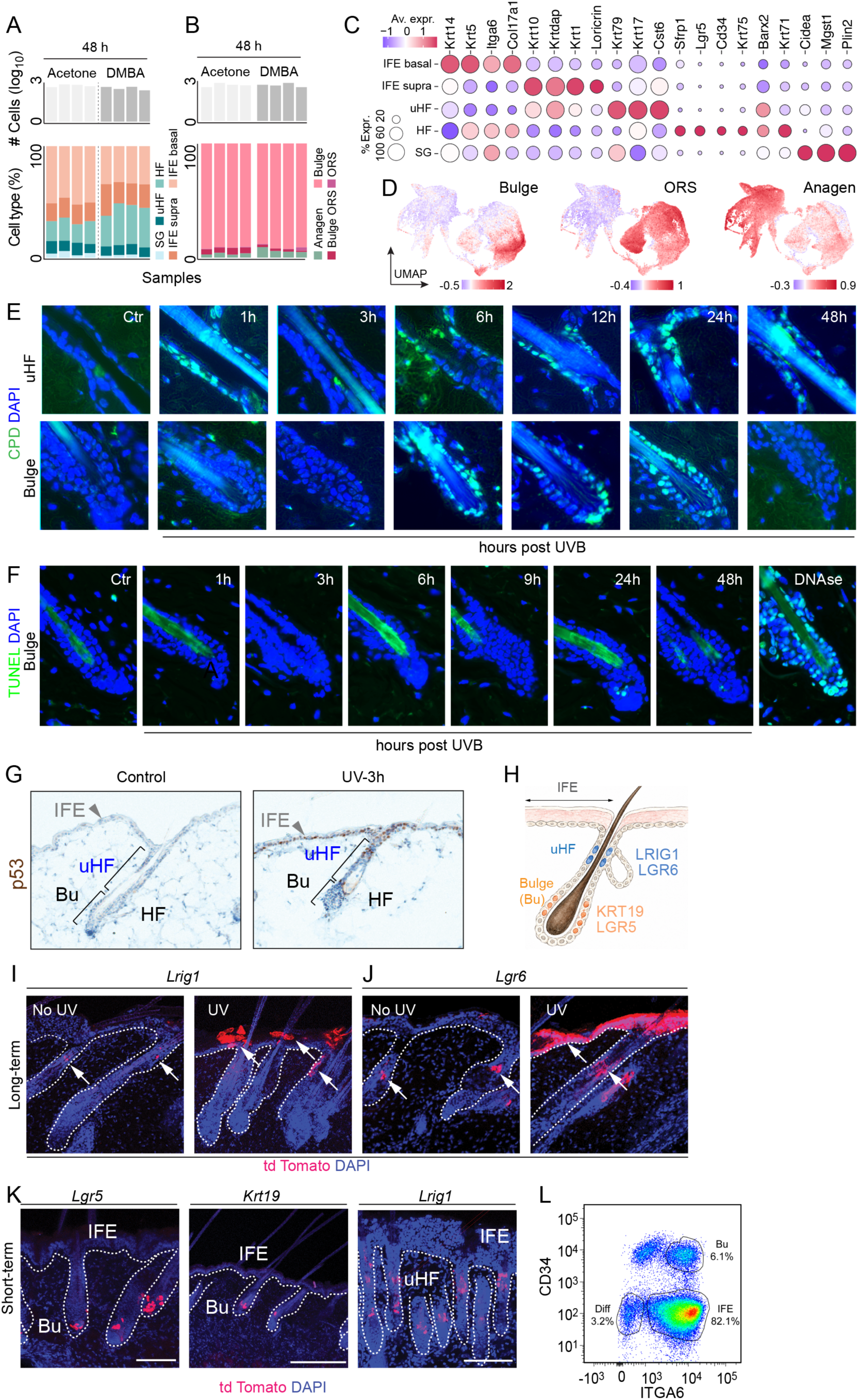
Hair follicle bulge stem cells show inherently different responses to mutagenic stress than interfollicular epidermal cells. (A, B) Number of cells (top) and proportion of cell types (bottom) of interfollicular (A) and follicular (B) sub-compartments. Each bar represents one animal. (C) Cell type-specific marker genes used to annotate the indicated sub-compartments. Dot size shows the fraction of detected cells. HF: Hair follicle. uHF: Upper hair follicle. ORS: Outer root sheath. IFE: Interfollicular epidermis. Supra: Suprabasal. SG: Sebaceous glands. (D) Module scores on a UMAP representing the average expression of genes within the defined gene sets (Methods) after subtraction of aggregated expression of control gene sets. (E, F) Detection of UV DNA lesion CPD (cyclopirimidine dimers) (E) and dead cells (TUNEL) (F) in control (Ctr) and UVB-treated skin. DNAse: Positive control for TUNEL staining. (H) Immunohistochemistry staining of P53-positive nuclei in the IFE upper hair follicle (uHF) in control or UVB-treated skin 3 hours (h) after exposure. (H) Illustration of epidermal populations analysed by lineage tracing. (I, J) Detection of tdTomato+ cells in untreated (No UV) and repeatedly UVB-irradiated back skin (3 x per week for a total of 3 weeks) in genetically labelled *Lrig*^+^ and *Lgr6*^+^ cell populations. Arrows indicate the location of tdTomato-positive cells. Dotted line: basement membrane. (K) Detection of tdTomato+ cells in back skin section 48 hours after exposure to a single dose of UVB in genetically labelled *Lgr5*^+^, *Krt19*^+^ and *Lrig1*^+^ cell populations. (L) Representative flow-cytometry gating strategy to sort viable bulge stem (CD34+/ITGA6+) and undifferentiated IFE (CD34-/ITGA6+) cells.

**Supplementary Figure 3.**
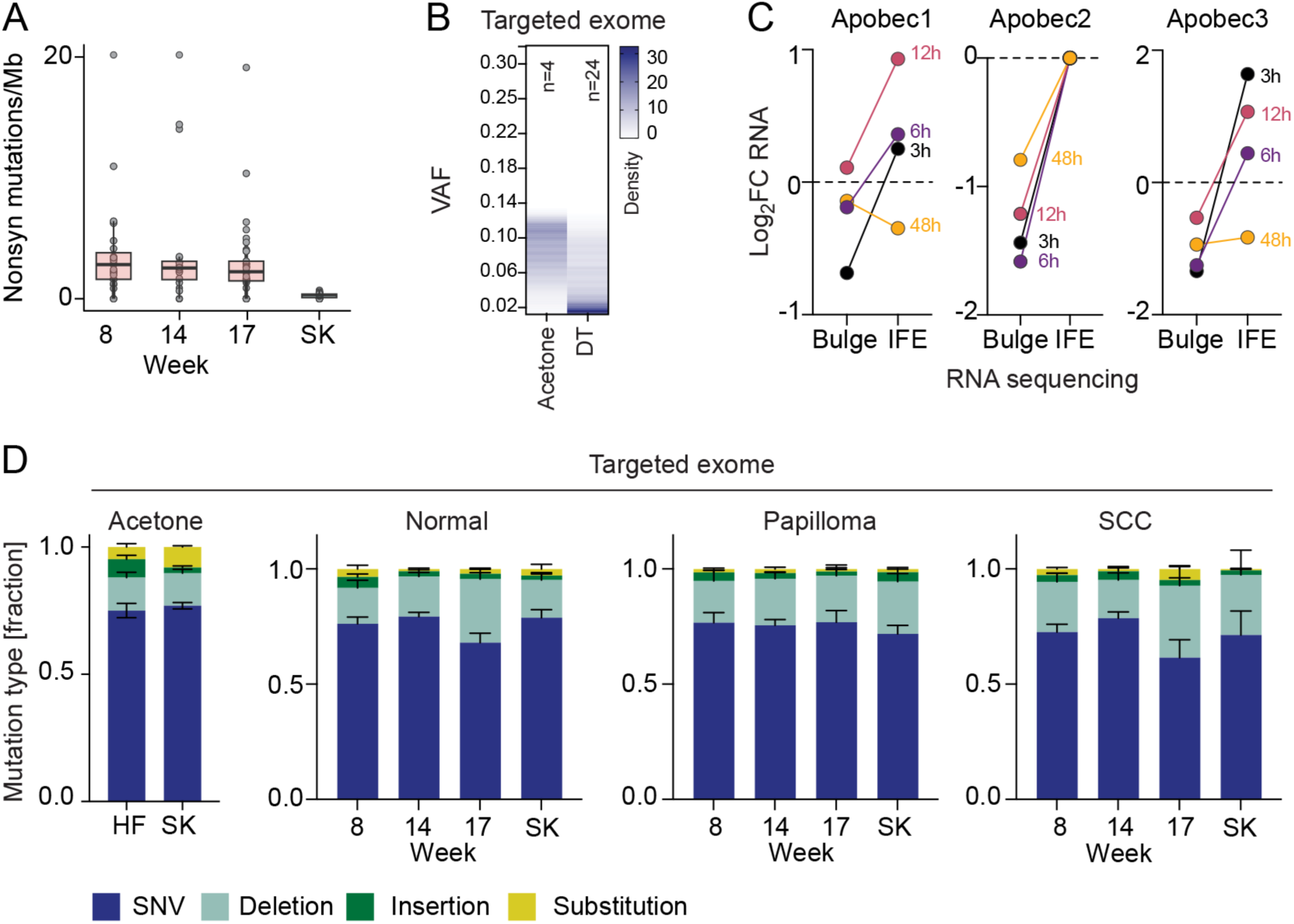
Mutation burden and signature in normal and tumour-prone epidermal regions. (**A**) Number of non-synonymous mutations per megabase (Mb) over time in hair follicles and in skin (SK). Points represent individual biopsies; boxplots show the median, interquartile range, and 1.5 IQR whiskers. (**B**) Heatmap showing density of VAF distribution in skin biopsies from acetone and DMBA/TPA (DT) conditions using targeted exome sequencing data. A beta-binomial model of ALT versus REF read counts, controlling for callable territory, was used to compare skin VAF distributions. Acetone skin mutations showed significantly higher allelic frequencies than mutations in skin samples of mice treated with DMBA/TPA (odds ratio = 1.81, BH-adjusted P = 0.0006). **(C)** Log_2_ fold change (FC) of APOBECs RNA expression in the bulge stem cells and interfollicular epidermis (IFE) at the indicated time points post-UVB exposure. Shown are all expressed APOBECs in skin. (**D**) Proportion of mutation types in hair follicles across time points (8, 14, 17 weeks) or skin biopsies (19 weeks) in normal or tumour-prone (papilloma, SCC) regions. Shown is mean ± SEM.

**Supplementary Figure 4.**
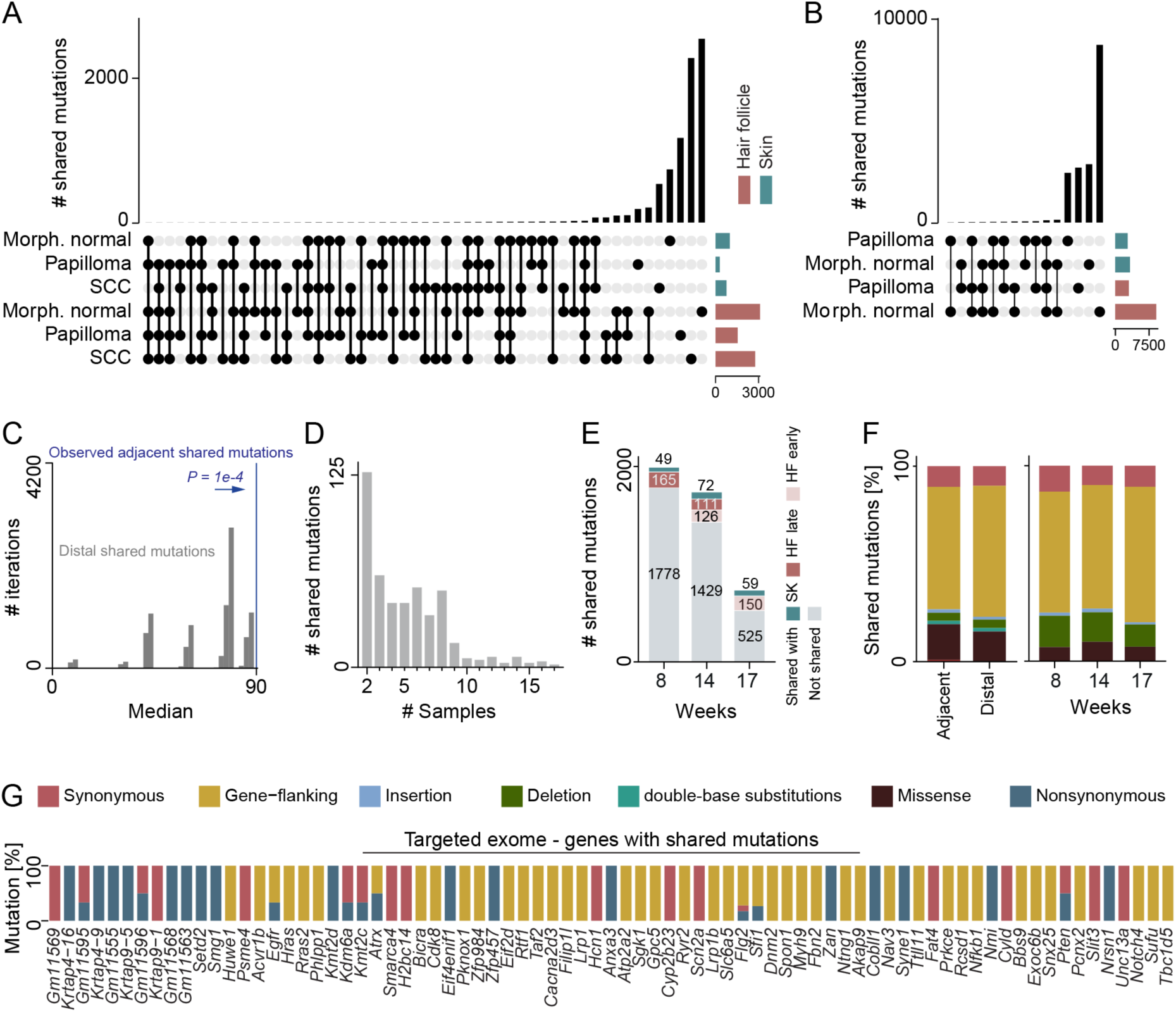
Dynamic changes of mutation burden during skin carcinogenesis. (**A, B**) Number of shared somatic mutations detected in hair follicles (HF) (green) and skin (brown) biopsies in macroscopically normal (normal) and tumour-destined regions (SCC, papilloma) obtained from targeted exome (A) or whole exome (B) sequencing. (**C**) Permutation analysis comparing the median observed number of shared mutations between spatially adjacent hair follicle and skin regions with the distribution obtained from randomly permuted pairings. Blue line: Median observed value. (**D**) Distribution showing the number of biopsies sharing somatic mutations identified by whole exome sequencing. (**E**) Number of mutations detected in hair follicles at weeks 8, 14, and 17 that were subsequently shared with endpoint skin biopsies or with later or earlier hair follicle time points (**F, G**) Distribution of shared genomic alterations identified by whole exome sequencing among adjacent and distal HF-skin (left) and in HF biopsies over the time course (right). Mutations are classified by the indicated colour shown in (G). (**G**) Complete mutation type distribution for genes that contain shared mutations.

**Supplementary Figure 5.**
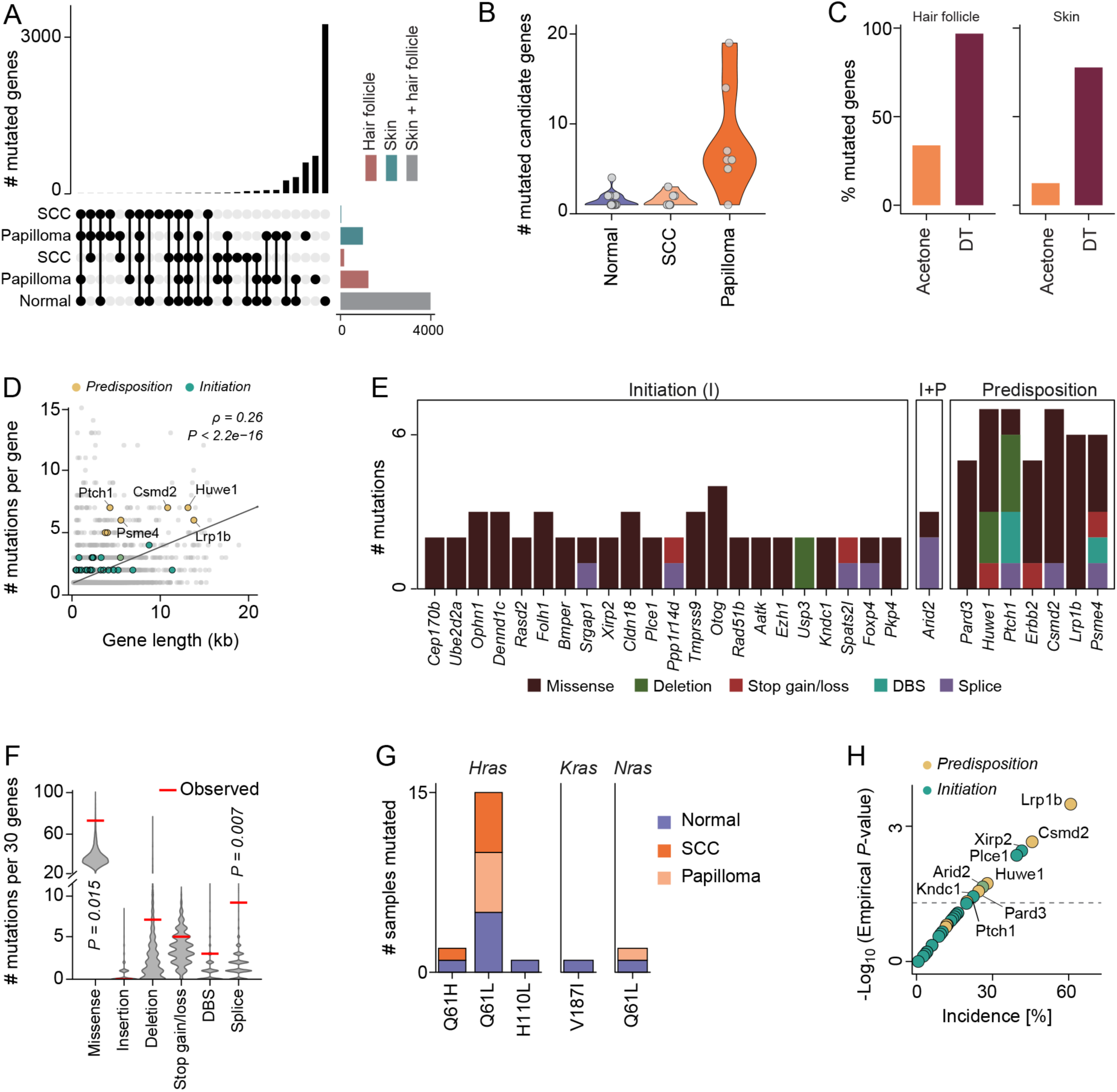
Longitudinal hair follicle profiling identifies candidate genes associated with tumour initiation and predisposition. **(A)** Number of recurrently mutated candidate genes shared between hair follicles (HF) and skin (SK) from macroscopically normal, papilloma, and squamous cell carcinoma (SCC) regions. **(B)** Number of mutated candidate genes in skin and hair follicles detected in macroscopically normal skin (Normal), papillomas, and SCCs. (**C**) Percentage of genes from the targeted panel (225 genes) harbouring at least one mutation, with a minimum of 10 informative reads and a VAF ≥ 0.01. (**D**) Correlation of gene length with the total number of detected mutations per gene. Each point represents one gene. Candidate genes are highlighted. Spearman’s correlation coefficient (ρ) and associated P-value are reported. (**E**) Number and type of nonsynonymous mutations identified in our candidate genes. Bars are coloured by mutation class (missense, deletion, stop gain/loss, double-base substitution (DBS), or splice-site mutation). (**F**) Distribution of mutation types across candidate genes. Violin plots show the distribution of mutation counts across 10,000 permuted sets of 30 genes for each mutation class (missense, insertion, deletion, stop gain/loss, double-base substitution (DBS), and splice-site mutation). Horizontal red bars: Observed number of mutations type for the 30 candidate genes. Empirical one-sided *P* values are shown for mutation types significantly enriched relative to the random distribution. **(G)** Number of samples carrying nonsynonymous mutations in *Hras*, *Kras*, and *Nras*, stratified by sample morphology. The known DMBA-associated *HRAS^Q61L^*mutation is detected in all three morphologies. (**H**) Correlation of human incidences of the candidate genes with enrichment significance (−log_10_ empirical P-value). The dashed line indicates the empirical significance threshold (P = 0.05). Selected recurrently mutated candidate genes are labelled.

## References

1 Que, S. K. T., Zwald, F. O. & Schmults, C. D. Cutaneous squamous cell carcinoma: Incidence, risk factors, diagnosis, and staging. J Am Acad Dermatol 78, 237–247 (2018). 10.1016/j.jaad.2017.08.059

2 Abel, E. L., Angel, J. M., Kiguchi, K. & DiGiovanni, J. Multi-stage chemical carcinogenesis in mouse skin: fundamentals and applications. Nat Protoc 4, 1350–1362 (2009). 10.1038/nprot.2009.120

3 Nassar, D., Latil, M., Boeckx, B., Lambrechts, D. & Blanpain, C. Genomic landscape of carcinogen-induced and genetically induced mouse skin squamous cell carcinoma. Nat Med 21, 946–954 (2015). 10.1038/nm.3878

4 Balmain, A., Ramsden, M., Bowden, G. T. & Smith, J. Activation of the mouse cellular Harvey-ras gene in chemically induced benign skin papillomas. Nature 307, 658–660 (1984). 10.1038/307658a0

5 Quintanilla, M., Brown, K., Ramsden, M. & Balmain, A. Carcinogen-specific mutation and amplification of Ha-ras during mouse skin carcinogenesis. Nature 322, 78–80 (1986). 10.1038/322078a0

6 Bizub, D., Wood, A. W. & Skalka, A. M. Mutagenesis of the Ha-ras oncogene in mouse skin tumors induced by polycyclic aromatic hydrocarbons. Proc Natl Acad Sci U S A 83, 6048–6052 (1986). 10.1073/pnas.83.16.6048

7 Su, F. et al. RAS mutations in cutaneous squamous-cell carcinomas in patients treated with BRAF inhibitors. N Engl J Med 366, 207–215 (2012). 10.1056/NEJMoa1105358

8 Lee, J. H. & Choi, S. Deciphering the molecular mechanisms of stem cell dynamics in hair follicle regeneration. Exp Mol Med 56, 110–117 (2024). 10.1038/s12276-023-01151-5

9 Derks, L. L. M. & van Boxtel, R. Stem cell mutations, associated cancer risk, and consequences for regenerative medicine. Cell stem cell 30, 1421–1433 (2023). 10.1016/j.stem.2023.09.008

10 Reeves, M. Q., Kandyba, E., Harris, S., Del Rosario, R. & Balmain, A. Multicolour lineage tracing reveals clonal dynamics of squamous carcinoma evolution from initiation to metastasis. Nat Cell Biol 20, 699–709 (2018). 10.1038/s41556-018-0109-0

11 Lapouge, G. et al. Identifying the cellular origin of squamous skin tumors. Proc Natl Acad Sci U S A 108, 7431–7436 (2011). 1012720108 [pii] 10.1073/pnas.1012720108

12 White, A. C. et al. Defining the origins of Ras/p53-mediated squamous cell carcinoma. Proc Natl Acad Sci U S A 108, 7425–7430 (2011). 1012670108 [pii] 10.1073/pnas.1012670108

13 Shendrik, I., Crowson, A. N. & Magro, C. M. Follicular cutaneous squamous cell carcinoma: an under-recognized neoplasm arising from hair appendage structures. Br J Dermatol 169, 384–388 (2013). 10.1111/bjd.12374

14 Winge, M. C. G. et al. Advances in cutaneous squamous cell carcinoma. Nat Rev Cancer 23, 430–449 (2023). 10.1038/s41568-023-00583-5

15 Greco, V. et al. A two-step mechanism for stem cell activation during hair regeneration. Cell stem cell 4, 155–169 (2009).

16 Faurschou, A., Haedersdal, M., Poulsen, T. & Wulf, H. C. Squamous cell carcinoma induced by ultraviolet radiation originates from cells of the hair follicle in mice. Exp Dermatol 16, 485–489 (2007). 10.1111/j.1600-0625.2007.00551.x

17 Mori, T. et al. Simultaneous establishment of monoclonal antibodies specific for either cyclobutane pyrimidine dimer or (6-4)photoproduct from the same mouse immunized with ultraviolet-irradiated DNA. Photochem Photobiol 54, 225–232 (1991). 10.1111/j.1751-1097.1991.tb02010.x

18 You, Y. H. et al. Cyclobutane pyrimidine dimers are responsible for the vast majority of mutations induced by UVB irradiation in mammalian cells. J Biol Chem 276, 44688–44694 (2001). 10.1074/jbc.M107696200

19 Chen, D. et al. Cell cycle duration determines oncogenic transformation capacity. Nature 641, 1309–1318 (2025). 10.1038/s41586-025-08935-x

20 Cotsarelis, G., Sun, T. T. & Lavker, R. M. Label-retaining cells reside in the bulge area of pilosebaceous unit: implications for follicular stem cells, hair cycle, and skin carcinogenesis. Cell 61, 1329–1337 (1990). 10.1016/0092-8674(90)90696-c

21 Braun, K. M. et al. Manipulation of stem cell proliferation and lineage commitment: visualisation of label-retaining cells in wholemounts of mouse epidermis. Development 130, 5241–5255 (2003).

22 Wilson, C. et al. Cells within the bulge region of mouse hair follicle transiently proliferate during early anagen: heterogeneity and functional differences of various hair cycles. Differentiation 55, 127–136 (1994). 10.1046/j.1432-0436.1994.5520127.x

23 Sachs, N. et al. Loss of integrin alpha3 prevents skin tumor formation by promoting epidermal turnover and depletion of slow-cycling cells. Proc Natl Acad Sci U S A 109, 21468–21473 (2012). 10.1073/pnas.1204614110

24 Ito, M. et al. Stem cells in the hair follicle bulge contribute to wound repair but not to homeostasis of the epidermis. Nat Med 11, 1351–1354 (2005).

25 Hameetman, L. et al. Molecular profiling of cutaneous squamous cell carcinomas and actinic keratoses from organ transplant recipients. BMC Cancer 13, 58 (2013). 10.1186/1471-2407-13-58

26 Yilmaz, A. S. et al. Differential mutation frequencies in metastatic cutaneous squamous cell carcinomas versus primary tumors. Cancer 123, 1184–1193 (2017). 10.1002/cncr.30459

27 Shea, L. K. et al. Combined Kdm6a and Trp53 Deficiency Drives the Development of Squamous Cell Skin Cancer in Mice. J Invest Dermatol 143, 232–241 e236 (2023). 10.1016/j.jid.2022.08.037

28 Yadav, R. et al. Atypical Site of Presentation of a Rare Type of SMARCA4-Positive Cutaneous Squamous Cell Carcinoma of the Skin: Case Report and Review of the Literature. J Investig Med High Impact Case Rep 12, 23247096241271977 (2024). 10.1177/23247096241271977

29 Alameda, J. P. et al. CYLD Inhibits the Development of Skin Squamous Cell Tumors in Immunocompetent Mice. Int J Mol Sci 22 (2021). 10.3390/ijms22136736

30 Mandemaker, I. K. et al. DNA damage-induced replication stress results in PA200-proteasome-mediated degradation of acetylated histones. EMBO Rep 19 (2018). 10.15252/embr.201745566

31 Ling, J. et al. RAS-mediated suppression of PAR3 and its effects on SCC initiation and tissue architecture occur independently of hyperplasia. J Cell Sci 133 (2020). 10.1242/jcs.249102

32 Koike, C. et al. Introduction of wild-type patched gene suppresses the oncogenic potential of human squamous cell carcinoma cell lines including A431. Oncogene 21, 2670–2678 (2002). 10.1038/sj.onc.1205370

33 Inoue, S. et al. Mule/Huwe1/Arf-BP1 suppresses Ras-driven tumorigenesis by preventing c-Myc/Miz1-mediated down-regulation of p21 and p15. Genes Dev 27, 1101–1114 (2013). 10.1101/gad.214577.113

34 Zhang, R. & Song, C. Loss of CSMD1 or 2 may contribute to the poor prognosis of colorectal cancer patients. Tumour Biol 35, 4419–4423 (2014). 10.1007/s13277-013-1581-6

35 Wadt, K. A. et al. Germline RAD51B truncating mutation in a family with cutaneous melanoma. Fam Cancer 14, 337–340 (2015). 10.1007/s10689-015-9781-4

36 Hu, X. et al. PLCE1 Polymorphisms Are Associated With Gastric Cancer Risk: The Changes in Protein Spatial Structure May Play a Potential Role. Frontiers in genetics 12, 714915 (2021). 10.3389/fgene.2021.714915

37 Li, X. et al. Distinct Subtypes of Gastric Cancer Defined by Molecular Characterization Include Novel Mutational Signatures with Prognostic Capability. Cancer Res 76, 1724–1732 (2016). 10.1158/0008-5472.CAN-15-2443

38 Yao, F. et al. Recurrent Fusion Genes in Gastric Cancer: CLDN18-ARHGAP26 Induces Loss of Epithelial Integrity. Cell reports 12, 272–285 (2015). 10.1016/j.celrep.2015.06.020

39 Hodis, E. et al. A landscape of driver mutations in melanoma. Cell 150, 251–263 (2012). 10.1016/j.cell.2012.06.024

40 Boudhraa, Z., Carmona, E., Provencher, D. & Mes-Masson, A. M. Ran GTPase: A Key Player in Tumor Progression and Metastasis. Front Cell Dev Biol 8, 345 (2020). 10.3389/fcell.2020.00345

41 Li, Y. Y. et al. Genomic analysis of metastatic cutaneous squamous cell carcinoma. Clin Cancer Res 21, 1447–1456 (2015). 10.1158/1078-0432.CCR-14-1773

42 Pickering, C. R. et al. Mutational landscape of aggressive cutaneous squamous cell carcinoma. Clin Cancer Res 20, 6582–6592 (2014). 10.1158/1078-0432.CCR-14-1768

43 Chang, D. & Shain, A. H. The landscape of driver mutations in cutaneous squamous cell carcinoma. NPJ Genom Med 6, 61 (2021). 10.1038/s41525-021-00226-4

44 Thind, A. S. et al. Whole genome analysis reveals the genomic complexity in metastatic cutaneous squamous cell carcinoma. Frontiers in oncology 12, 919118 (2022). 10.3389/fonc.2022.919118

45 Guo, Y. et al. Clinical significance of the correlation between PLCE 1 and PRKCA in esophageal inflammation and esophageal carcinoma. Oncotarget 8, 33285–33299 (2017). 10.18632/oncotarget.16635

46 Kandyba, E. et al. Chemically induced skin tumors arise from long-lived stem cells of the upper hair follicle. *Science*, eadv8291 (2026). 10.1126/science.adv8291

47 Beerman, I., Seita, J., Inlay, M. A., Weissman, I. L. & Rossi, D. J. Quiescent hematopoietic stem cells accumulate DNA damage during aging that is repaired upon entry into cell cycle. Cell stem cell 15, 37–50 (2014). 10.1016/j.stem.2014.04.016

48 Murai, K. et al. Epidermal Tissue Adapts to Restrain Progenitors Carrying Clonal p53 Mutations. Cell stem cell 23, 687–699 e688 (2018). 10.1016/j.stem.2018.08.017

49 Martincorena, I. et al. Tumor evolution. High burden and pervasive positive selection of somatic mutations in normal human skin. Science 348, 880–886 (2015). 10.1126/science.aaa6806

50 Madisen, L. et al. A robust and high-throughput Cre reporting and characterization system for the whole mouse brain. Nat Neurosci 13, 133–140 (2010). 10.1038/nn.2467

51 Means, A. L., Xu, Y., Zhao, A., Ray, K. C. & Gu, G. A CK19(CreERT) knockin mouse line allows for conditional DNA recombination in epithelial cells in multiple endodermal organs. Genesis 46, 318–323 (2008). 10.1002/dvg.20397

52 Jaks, V. et al. Lgr5 marks cycling, yet long-lived, hair follicle stem cells. Nat Genet 40, 1291–1299 (2008). ng.239 [pii] 10.1038/ng.239

53 Page, M. E., Lombard, P., Ng, F., Gottgens, B. & Jensen, K. B. The epidermis comprises autonomous compartments maintained by distinct stem cell populations. Cell stem cell 13, 471–482 (2013). 10.1016/j.stem.2013.07.010

54 Snippert, H. J. et al. Lgr6 marks stem cells in the hair follicle that generate all cell lineages of the skin. Science 327, 1385–1389 (2010). 327/5971/1385 [pii] 10.1126/science.1184733

55 Jensen, K. B., Driskell, R. R. & Watt, F. M. Assaying proliferation and differentiation capacity of stem cells using disaggregated adult mouse epidermis. Nat Protoc 5, 898–911 (2010). nprot.2010.39 [pii] 10.1038/nprot.2010.39

56 Blanco, S. et al. Stem cell function and stress response are controlled by protein synthesis. Nature 534, 335–340 (2016). 10.1038/nature18282

57 Blanco, S. et al. The RNA-methyltransferase Misu (NSun2) poises epidermal stem cells to differentiate. PLoS Genet 7, e1002403 (2011). 10.1371/journal.pgen.1002403 PGENETICS-D-11-00619 [pii]

58. Schindelin, J., et al. Fiji: an open-source platform for biological-image analysis. Nat Methods 9, 676-682 (2012). 10.1038/nmeth.2019

59 Reisinger, E. et al. OTP: An automatized system for managing and processing NGS data. J Biotechnol 261, 53–62 (2017). 10.1016/j.jbiotec.2017.08.006

60 Dobin, A. et al. STAR: ultrafast universal RNA-seq aligner. Bioinformatics 29, 15–21 (2013). 10.1093/bioinformatics/bts635

61 Joost, S. et al. The Molecular Anatomy of Mouse Skin during Hair Growth and Rest. Cell stem cell 26, 441–457 e447 (2020). 10.1016/j.stem.2020.01.012

62 Love, M. I., Huber, W. & Anders, S. Moderated estimation of fold change and dispersion for RNA-seq data with DESeq2. Genome Biol 15, 550 (2014). 10.1186/s13059-014-0550-8

63 Korotkevich G., S. V., Budin N., Shpak B., Artyomov M.N., Sergushichev A. Fast gene set enrichment analysis. bioRxiv (2021). 10.1101/060012

64 Subramanian, A. et al. Gene set enrichment analysis: a knowledge-based approach for interpreting genome-wide expression profiles. Proc Natl Acad Sci U S A 102, 15545–15550 (2005). 10.1073/pnas.0506580102

65 Liberzon, A. et al. The Molecular Signatures Database (MSigDB) hallmark gene set collection. Cell Syst 1, 417–425 (2015). 10.1016/j.cels.2015.12.004

66 Castanza, A. S. et al. Extending support for mouse data in the Molecular Signatures Database (MSigDB). Nat Methods 20, 1619–1620 (2023). 10.1038/s41592-023-02014-7

67 Cheng, X., Yan, J., Liu, Y., Wang, J. & Taubert, S. eVITTA: a web-based visualization and inference toolbox for transcriptome analysis. Nucleic Acids Res 49, W207–W215 (2021). 10.1093/nar/gkab366

